# Exome-by-phenome-wide rare variant gene burden association with electronic health record phenotypes

**DOI:** 10.1101/798330

**Authors:** Joseph Park, Nathan Katz, Xinyuan Zhang, Anastasia M Lucas, Anurag Verma, Renae L Judy, Rachel L Kember, Regeneron Genetics Center, Jinbo Chen, Scott M Damrauer, Marylyn D Ritchie, Daniel J Rader

**Author notes:** To whom correspondence should be addressed: Daniel J. Rader, MD, Perelman School of Medicine at the University of Pennsylvania and Children’s Hospital of Philadelphia, 11-125 Smilow Center for Translational Research, 3400 Civic Center Blvd, Philadelphia, PA 19104-5158.

## Abstract

**Background:** By coupling large-scale DNA sequencing with electronic health records (EHR), “genome-first” approaches can enhance our understanding of the contribution of rare genetic variants to disease. Aggregating rare, loss-of-function variants in a candidate gene into a “gene burden” to test for association with EHR phenotypes can identify both known and novel clinical implications for the gene in human disease. However, this methodology has not yet been applied on both an exome-wide and phenome-wide scale, and the clinical ontologies of rare loss-of-function variants in many genes have yet to be described.

**Methods:** We leveraged whole exome sequencing (WES) data in participants (N=11,451) in the Penn Medicine Biobank (PMBB) to address on an exome-wide scale the association of a burden of rare loss-of-function variants in each gene with diverse EHR phenotypes using a phenome-wide association study (PheWAS) approach. For discovery, we collapsed rare (minor allele frequency (MAF) ≤ 0.1%) predicted loss-of-function (pLOF) variants (*i.e.* frameshift insertions/deletions, gain/loss of stop codon, or splice site disruption) per gene to perform a gene burden PheWAS. Subsequent evaluation of the significant gene burden associations was done by collapsing rare (MAF ≤ 0.1%) missense variants with Rare Exonic Variant Ensemble Learner (REVEL) scores ≥ 0.5 into corresponding yet distinct gene burdens, as well as interrogation of individual low-frequency to common (MAF > 0.1%) pLOF variants and missense variants with REVEL≥ 0.5. We replicated our findings using the UK Biobank’s (UKBB) whole exome sequence dataset (N=49,960).

**Results:** From the pLOF-based discovery phase, we identified 106 gene burdens with phenotype associations at p<10^-6^ from our exome-by-phenome-wide association studies. Positive-control associations included *TTN* (cardiomyopathy, p=7.83E-13), *MYBPC3* (hypertrophic cardiomyopathy, p=3.48E-15), *CFTR* (cystic fibrosis, p=1.05E-15), *CYP2D6* (adverse effects due to opiates/narcotics, p=1.50E-09), and *BRCA2* (breast cancer, p=1.36E-07). Of the 106 genes, 12 gene-phenotype relationships were also detected by REVEL-informed missense-based gene burdens and 19 by single-variant analyses, demonstrating the robustness of these gene-phenotype relationships. Three genes showed evidence of association using both additional methods (*BRCA1*, *CFTR*, *TGM6*), leading to a total of 28 robust gene-phenotype associations within PMBB. Furthermore, replication studies in UKBB validated 30 of 106 gene burden associations, of which 12 demonstrated robustness in PMBB.

**Conclusion:** Our study presents 12 exome-by-phenome-wide robust gene-phenotype associations, which include three proof-of-concept associations and nine novel findings. We show the value of aggregating rare pLOF variants into gene burdens on an exome-wide scale for unbiased association with EHR phenotypes to identify novel clinical ontologies of human genes. Furthermore, we show the significance of evaluating gene burden associations through complementary, yet non-overlapping genetic association studies from the same dataset. Our results suggest that this approach applied to even larger cohorts of individuals with WES or whole-genome sequencing data linked to EHR phenotype data will yield many new insights into the relationship of genetic variation and disease phenotypes.

## Introduction

A “genome-first” approach, in which genetic variants of interest are identified and then subsequently associated with phenotypes, has the potential to inform the genetic basis of human phenotypes and disease.^1, 2^ This approach can be applied to healthcare populations with extensive electronic health record (EHR) phenotype data, thus permitting “phenome-wide association studies” (PheWAS) as an unbiased approach to determining the clinical impact of specific genetic variants.^3–5^ Genome-first approaches like PheWAS have thus far been characterized primarily by univariate analyses focusing on individual common variants. Rare coding variants are predicted to have larger effect sizes on phenotype and would be of interest to study in a similar unbiased manner, but are too underpowered to study in a univariate fashion.^3^ Statistical aggregation tests that interrogate the cumulative effects of multiple rare variants in a gene (*i.e.* ‘gene burden’) not only increase the statistical power of regression analyses, but also enable gene-based association studies to describe the implications of loss of gene function in human disease.^6^ Previously, we leveraged the Penn Medicine Biobank (PMBB, University of Pennsylvania), a large academic biobank with whole exome sequencing (WES) data linked to EHR data, to show that aggregating rare, loss-of-function variants in a single gene or targeted sets of genes to conduct gene burden PheWAS has the potential to uncover novel pleiotropic relationships between the gene and human disease.^7–9^

This approach has not yet been applied on an exome-wide scale, and the clinical ontologies of loss-of-function variants in many genes have yet to be described. Thus, we applied gene burden PheWAS on an exome-wide scale utilizing WES data to conduct exome-by-phenome-wide association studies (ExoPheWAS) to evaluate in detail the phenotypes (*i.e.* Phecodes)^10^ associated with rare predicted loss-of-function (pLOF) variants on a gene-by-gene basis across the human exome. We further evaluated the robustness of the gene-disease associations identified via our pLOF-based gene burden discovery analyses by testing the associations between other likely deleterious exonic variants and the significant disease phenotypes. We also replicated findings in the UK Biobank (UKBB) WES dataset. Furthermore, we interrogated the EHR data for quantitative phenotypic traits via analyses of clinical imaging and laboratory measurements. Our findings represent the first report combining a genome-first approach to examining the clinical effects of pLOF variants per gene on an exome-wide level with subsequent validation through interrogation of other variable regions in the same genes.

## Materials and Methods

### Setting and study participants

All individuals who were recruited for the Penn Medicine Biobank (PMBB) are patients of clinical practice sites of the University of Pennsylvania Health System. Appropriate consent was obtained from each participant regarding storage of biological specimens, genetic sequencing, access to all available electronic health record (EHR) data, and permission to recontact for future studies.

In addition to our robustness validation analyses within PMBB, replication analyses were conducted using the WES dataset from the UK Biobank (UKBB) for evaluation of the robustness of gene-phenotype associations identified in PMBB.

### Exome sequencing

This PMBB study dataset included a subset of 11,451 individuals in the PMBB who have undergone whole-exome sequencing (WES). For each individual, we extracted DNA from stored buffy coats and then obtained exome sequences as generated by the Regeneron Genetics Center (Tarrytown, NY). These sequences were mapped to GRCh37 as previously described.^7^ Furthermore, for subsequent phenotypic analyses, we removed samples with low exome sequencing coverage (*i.e.* less than 75% of targeted bases achieving 20x coverage), high missingness (*i.e.* greater than 5% of targeted bases), high heterozygosity, dissimilar reported and genetically determined sex, genetic evidence of sample duplication, and cryptic relatedness (*i.e.* closer than 3^rd^ degree relatives), leading to a total of 10,996 individuals.

For replication studies in UKBB, we interrogated the 42,228 individuals with ICD-10 diagnosis codes available among the 49,960 individuals who had WES data as generated by the Functional Equivalence (FE) pipeline. The PLINK files for exome sequencing provided by UKBB were based on mappings to GRCh38. Access to the UK Biobank for this project was from Application 32133.

### Variant annotation and selection for association testing

For both PMBB and UKBB, genetic variants were annotated using ANNOVAR^11^ as predicted loss-of-function (pLOF) or missense variants according to the NCBI Reference Sequence (RefSeq) database.^12^ pLOF variants were defined as frameshift insertions/deletions, gain/loss of stop codon, or disruption of canonical splice site dinucleotides. Predicted deleterious missense variants were defined as those with Rare Exonic Variant Ensemble Learner (REVEL)^13^ scores ≥ 0.5. Minor allele frequencies for each variant were determined per Non-Finnish European and African/African-American minor allele frequencies reported by the Genome Aggregation Database (gnomAD)^14^.

### Clinical data collection

International Classification of Diseases Ninth Revision (ICD-9) and Tenth Revision (ICD-10) disease diagnosis codes and procedural billing codes, medications, and clinical imaging and laboratory measurements were extracted from the patients’ electronic health records (EHR) for PMBB. ICD-10 encounter diagnoses were mapped to ICD-9 via the Center for Medicare and Medicaid Services 2017 General Equivalency Mappings (https://www.cms.gov/Medicare/Coding/ICD10/2017-ICD-10-CM-and-GEMs.html) and manual curation. Phenotypes for each individual were then determined by mapping ICD-9 codes to distinct disease entities (*i.e.* Phecodes) via Phecode Map 1.2 using the R package “PheWAS”.^10, 15^ Patients were determined to have a certain disease phenotype if they had the corresponding ICD diagnosis on two or more dates, while phenotypic controls consisted of individuals who never had the ICD code. Individuals with an ICD diagnosis on only one date as well as individuals under control exclusion criteria based on PheWAS phenotype mapping protocols were not considered in statistical analyses.

All laboratory values measured in the outpatient setting were extracted for participants from the time of enrollment in the Biobank until March 20, 2019; all units were converted to their respective clinical Traditional Units. Minimum, median, and maximum measurements of each laboratory measurement were recorded for each individual and used for all association analyses. Inpatient and outpatient echocardiography measurements were extracted if available for participants from January 1, 2010 until September 9, 2016; outliers for each echocardiographic parameter (less than Q1 - 1.5*IQR or greater than Q3 + 1.5*IQR) were removed. Similarly, minimum, median, and maximum values for each parameter were recorded for each patient and used for association analyses. Additionally, in UKBB, we used the provided ICD-10 disease diagnosis codes for replication studies.

### Association studies

A phenome-wide association study (PheWAS) approach was used to determine the phenotypes associated with rare pLOF variants carried by individuals in PMBB for the discovery experiment.^10^ Each disease phenotype was tested for association with each gene burden or single variant using a logistic regression model adjusted for age, age^2^, gender, and the first ten principal components of genetic ancestry. We used an additive genetic model to collapse variants per gene via an extension of the fixed threshold approach.^16^ Given the high percentage of individuals of African ancestry present in PMBB, association analyses were performed separately in European and African genetic ancestries and combined with inverse variance weighted meta-analysis. Our association analyses considered only disease phenotypes with at least 20 cases, leading to the interrogation of 1000 total Phecodes. All association analyses were completed using R version 3.3.1 (Vienna, Austria).

We further evaluated the robustness of our gene-phenotype associations in PMBB by 1) associating the aggregation of rare predicted missense variants in a gene burden test and 2) testing pLOFs and predicted missense in a univariate association test across significant genes for a hypothesis-driven evaluation with the specific phenotype identified in the pLOF-based gene burden association as well as other related phenotypes in their corresponding Phecode families (integer part of Phecode). For example, to replicate an association of a gene burden with hypothetical Phecode 123.45, we associated other variants in the same gene with PheCode 123.45 as well as other related phenotypes under the Phecode family 123 (*e.g.* 123.6). Notably, we checked for the presence of mutual carriers between each gene’s pLOF-based gene burdens and subsequently interrogated missense-based gene burdens or single variants due to linkage disequilibrium and/or rare chance, and only reported replications for which the significant phenotypes’ associations were not being driven by mutual carriers. All association studies in PMBB were based on a logistic regression model adjusted for age, age^2^, gender, and the first ten principal components of genetic ancestry

Additionally, we replicated our findings in UKBB for genes of interest using pLOF-based gene burden, REVEL-informed missense-based gene burden, and/or univariate association analyses in PMBB. Phenotypes for each individual in UKBB with WES data was determined by mapping ICD-10 codes to Phecodes via Phecode Map 1.2b1 using the R package “PheWAS”.^15^ Patients were determined to have a certain disease phenotype if they had one or more encounters for the corresponding ICD diagnosis given the lack of individuals with more than two encounters per diagnosis, while phenotypic controls consisted of individuals who never had the ICD code. Individuals under control exclusion criteria based on PheWAS phenotype mapping protocols were not considered in statistical analyses. We conducted our replication association studies among the 34,629 individuals of European ancestry based on UKBB’s reported genetic ancestry grouping. Association statistics were calculated similarly to PMBB, such that each disease phenotype was tested for association with each gene burden or single variant using a logistic regression model adjusted for age, age^2^, gender, and the first ten principal components of genetic ancestry. All association analyses for UKBB were completed using R version 3.6.1 (Vienna, Austria).

### Statistical analyses of clinical measurements

In order to compare available measurements for echocardiographic parameters and serum laboratory values between carriers of predicted deleterious variants and genotypic controls, we utilized linear regression adjusted for age, age^2^, gender, and the first ten principal components of genetic ancestry in both the overall population and individuals of European ancestry alone. These analyses were conducted with the minimum, median, and maximum value as the dependent variable for each echocardiographic parameter and clinical lab measure. All statistical analyses, including PheWAS, were completed using R version 3.3.1 or R 3.6.1 (Vienna, Austria).

## Results

### Discovery: exome-by-phenome-wide gene burden analyses of rare pLOF variants

We interrogated our dataset of individuals with WES data in PMBB (Table 1) for carriers of rare (MAF ≤ 0.1%) pLOF variants. Assuming an additive disease model, the distribution of number of carriers for rare pLOF variants per gene was on a negative exponential (Figure S1). We chose to interrogate genes with at least 25 heterozygous carriers for rare pLOFs (N=1518 genes). We collapsed rare pLOF variants across these 1518 genes for exome-by-phenome-wide gene burden association analyses with 1000 binary EHR-derived diagnosis phenotypes (Figure 1). Given that p values for gene burden association studies interrogating rare protein truncating variants are inflated,^17^ we found that our associations roughly deviated from the fitted expected distribution at an observed p<E-06 (Figure S2). We identified 139 gene-disease relationships with p<E-06 across 106 unique gene burdens (Figure 2, Table S1). Among these, known positive-control associations included *TTN* with cardiomyopathy and related cardiac conduction disorders, *MYBPC3* with hypertrophic cardiomyopathy, *CFTR* with cystic fibrosis and bronchiectasis, *CYP2D6* with adverse effects due to opiates/narcotics, and *BRCA2* with breast cancer. We addressed potential issues regarding small sample sizes by using Firth’s penalized likelihood approach, and found that beta and significance estimates were consistent with exact logistic regression (Table S1).

**Figure 1.**
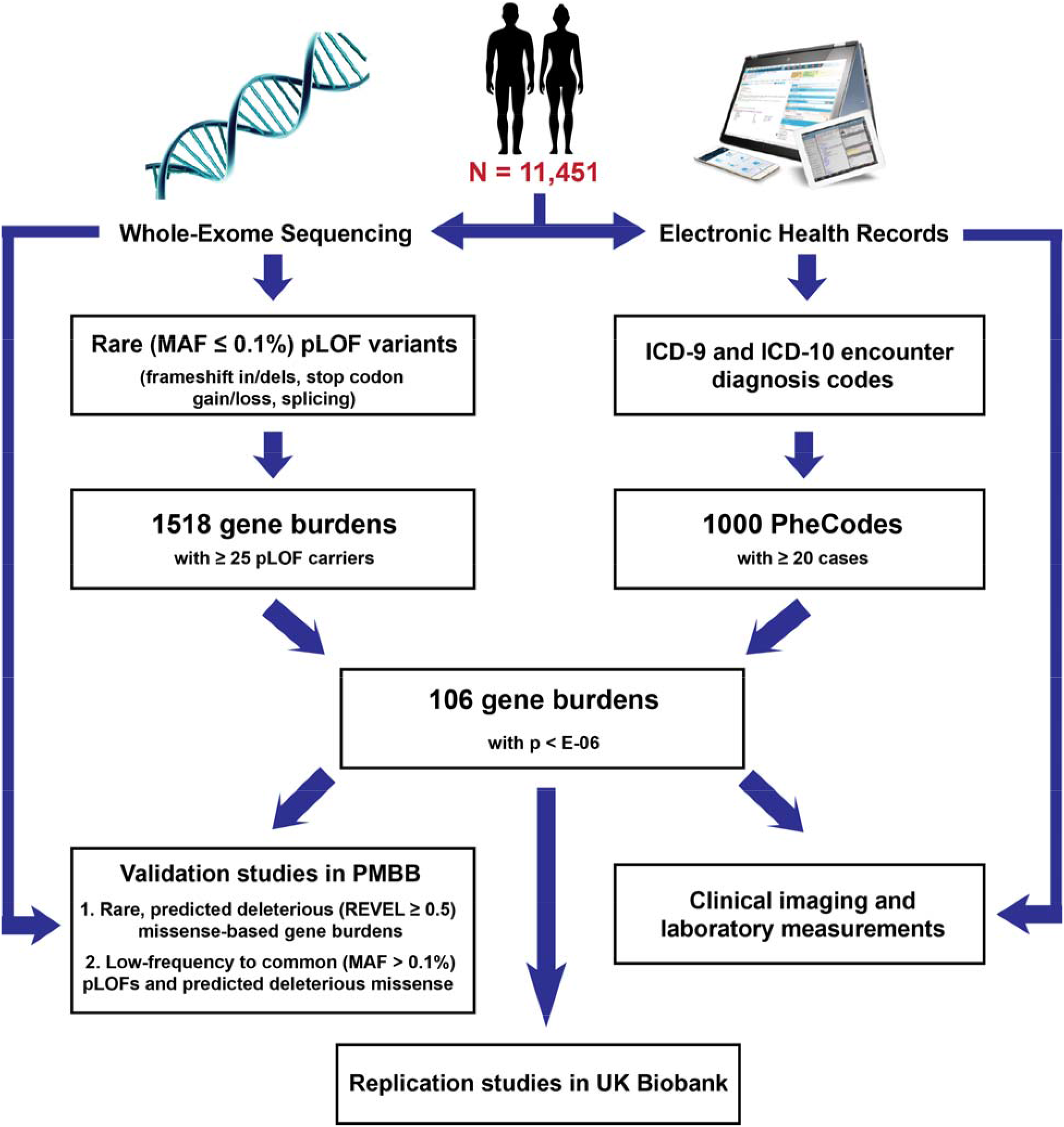
Description of methods for exome-by-phenome-wide association analyses using electronic health record phenotypes. Flowchart diagram outlining the major methodologies used for conducting our exome-by-phenome-wide association studies and evaluation of the robustness of association for major findings (*i.e.* 106 genes with associations p<E-06). The pathways starting with short descending arrows represent the discovery phase focusing on predicted loss-of-function (pLOF)-based gene burden association studies on an exome-by-phenome-wide scale in Penn Medicine Biobank (PMBB). “Validation studies in PMBB” represents evaluation of the robustness of gene-phenotype associations using REVEL-informed missense-based gene burdens and univariate analyses within PMBB. “Replication studies in UK Biobank” represents evaluation of the robustness of gene-phenotype associations using pLOF-based gene burdens, REVEL-informed missense-based gene burdens, and univariate analyses in UK Biobank.

**Figure 2.**
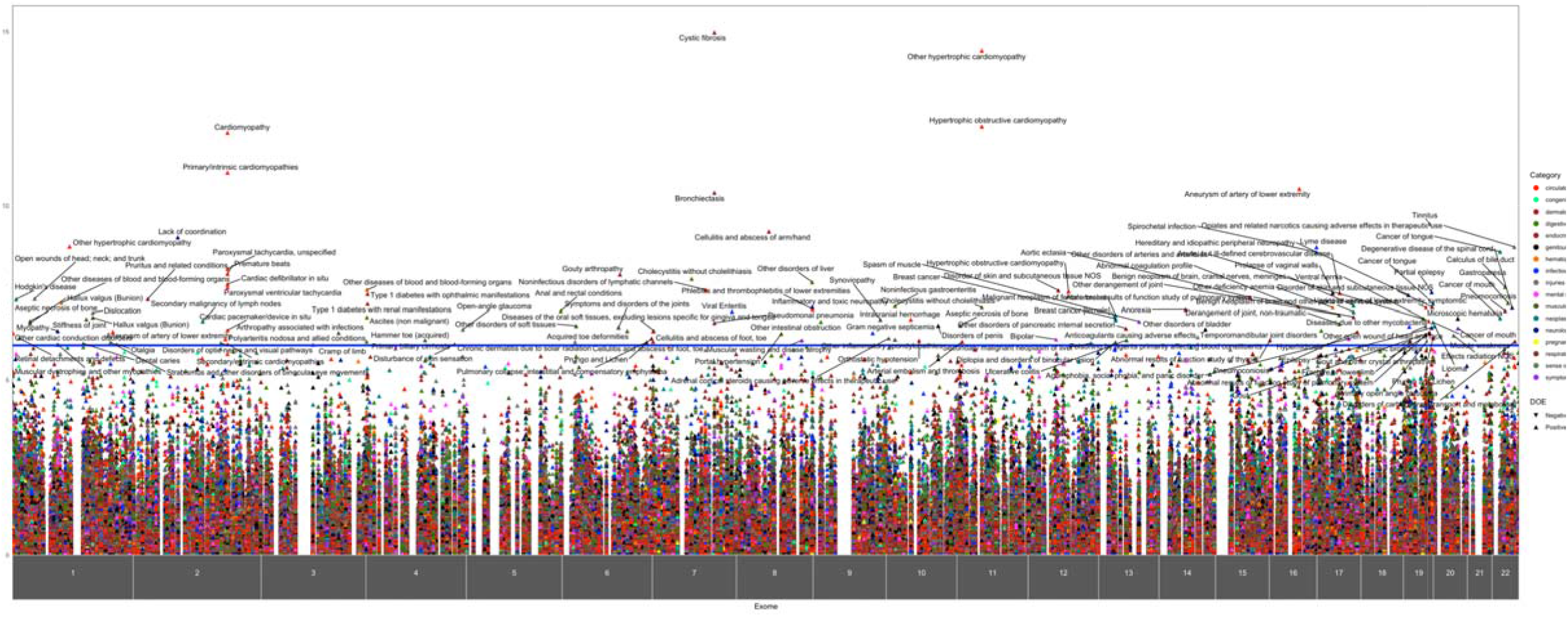
ExoPheWAS plot for Penn Medicine Biobank’s whole exome-sequenced cohort. Exome-by-Phenome-Wide Association Study (ExoPheWAS) plot of 1518 gene burdens collapsing rare (MAF ≤ 0.1%) predicted loss-of-function (pLOF) variants in Penn Medicine Biobank. The x-axis represents the exome and is organized by chromosomal location. Each gene’s associations are located along the x-axis according to the gene’s genomic locations per GRCh37. Each gene burden’s associations with 1000 Phecodes are plotted vertically above each gene, with the height of each point representing the –log_10_(p) of the association between the gene burden and Phecode. Each Phecode point is color-coded per Phecode group, and direction of effect is represented by the directionality of each triangular point. The blue line represents the significance threshold at p=E-06.

**Table 1:** Demographics of Penn Medicine Biobank’s whole exome-sequenced cohort. Demographic information and basic clinical diagnosis counts for the individuals with exome sequencing linked to electronic health records in Penn Medicine Biobank. Data is represented as count data followed by percent prevalence in the population where appropriate.

|  |  |
| --- | --- |
| **Basic demographics** |  |
| N | 10996 |
| Male, N (%) | 6527 (59.4%) |
| Median Age (at biobank entry), yr | 67.9 |
| **Race** |  |
| AFR, N (%) | 2203 (20.0) |
| AMR, N (%) | 307 (2.8) |
| EAS, N (%) | 79 (0.7) |
| EUR, N (%) | 8259 (75.1) |
| SAS, N (%) | 115 (1.0) |
| **Clinical Diagnoses** |  |
| Diabetes Mellitus, N (%) | 3534 (32.1) |
| Hypertension, N (%) | 8008 (72.8) |
| Coronary Artery Disease, N (%) | 4797 (43.6) |
| Myocardial Infarction, N (%) | 2228 (20.3) |
| Heart Failure, N (%) | 4200 (38.2) |

### Robustness evaluation via gene burden analyses of rare predicted deleterious missense variants

To evaluate the robustness of the gene-disease relationships discovered from our pLOF-based gene burden discovery analyses, we separately collapsed rare (MAF ≤ 0.1%), predicted deleterious (*i.e.* REVEL score of at least 0.5) missense variants (with no overlap with the pLOFs) across the 106 significant genes for hypothesis-driven associations with their corresponding phenotypes as well as other related phenotypes in their corresponding Phecode families (Figure 1). Of the 106 genes identified from the pLOF-based gene burden discovery analyses, 66 genes had at least five carriers for rare, predicted deleterious missense variants in PMBB (Table S2). Of these, 11 were significantly associated in the REVEL-informed missense-based gene burdens (Table 2, Table S3), namely *BBS10*, *BRCA1*, *CFTR*, *CILP, CYP2D6*, *MYBPC3*, *MYCBP2*, *PPP1R13L*, *RGS12*, *SCRN3*, and *TGM6,* and another two were borderline [*AIM1* (p=0.0501) and *POLN* (p=0.0553)].

**Table 2.**
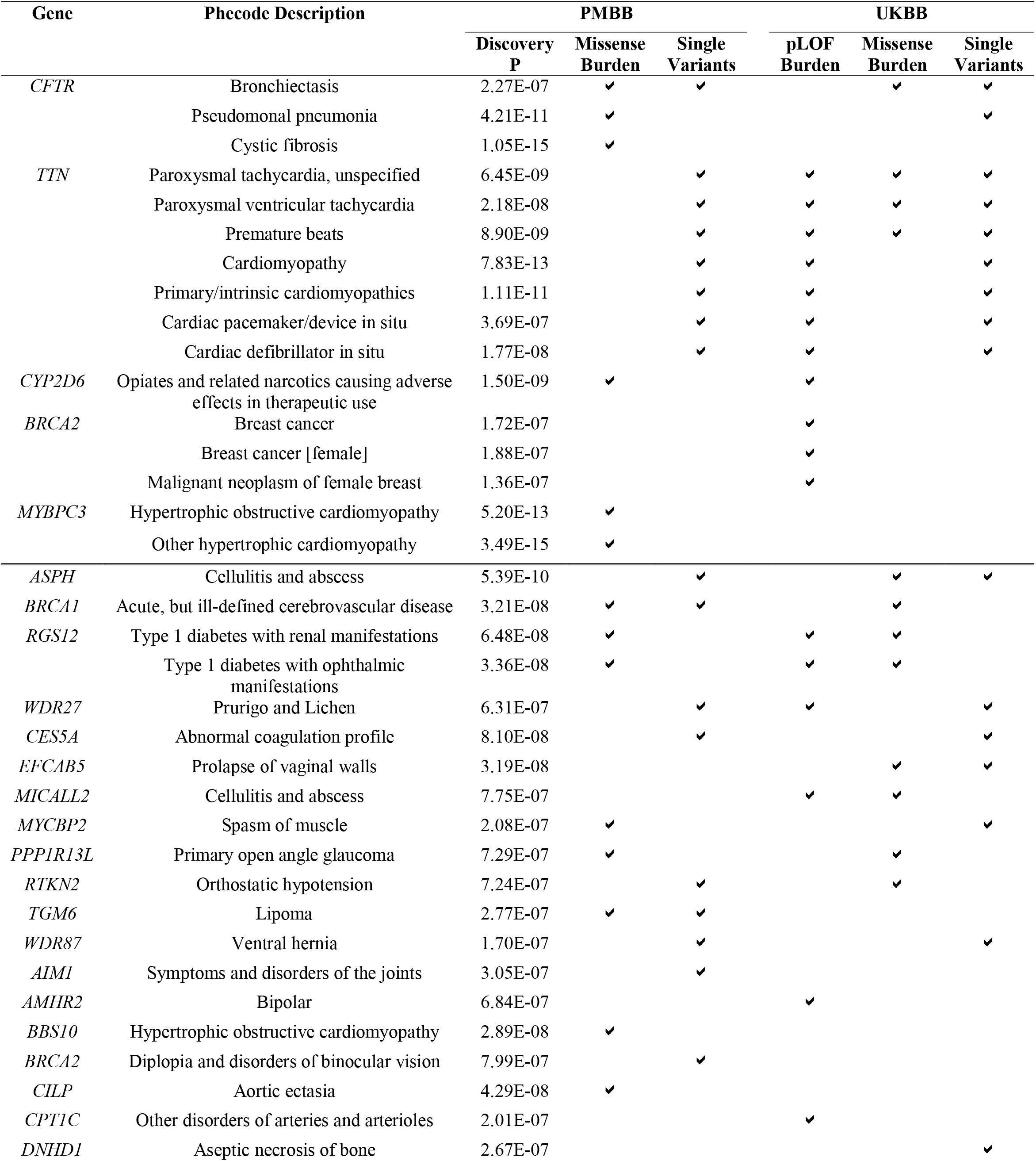
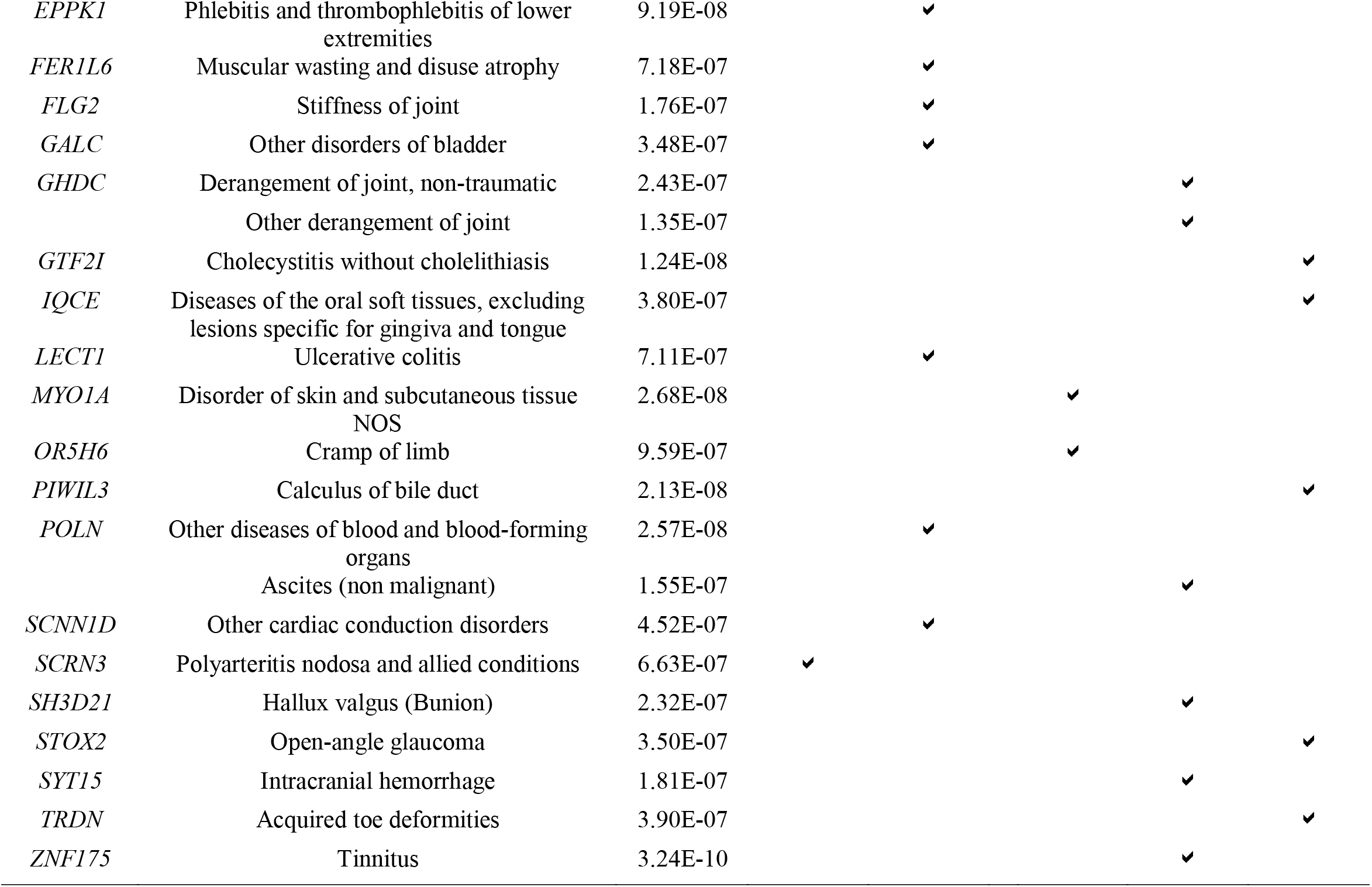
List of successfully evaluated gene-phenotype relationships in Penn Medicine Biobank and/or UK Biobank. List of genes among 106 gene burdens with phenotype associations at p<E-06 that were successfully evaluated for the robustness of their gene-phenotype relationships by interrogating REVEL-informed missense-based gene burdens and single variants in Penn Medicine Biobank (PMBB) and/or interrogating pLOF-based gene burdens, REVEL-informed missense-based gene burdens, and single variants in UK Biobank (UKBB). Each association is labeled with the corresponding p value from the discovery phase in PMBB, and checkmarks for successful evaluation for the respective validation/replication studies in PMBB/UKBB. Positive-control associations are listed on the top and are separated from novel associations by a double line. Novel associations are ranked by number of successful validation/replication studies in PMBB/UKBB, and then alphabetically by gene name. Associations are grouped by gene when possible.

### Robustness evaluation via univariate PheWAS of predicted deleterious variants

We also interrogated single variants in the 106 genes identified on discovery that were of sufficient frequency (MAF > 0.1%) and were not included in the gene burden analyses, including both pLOF variants and predicted deleterious (REVEL ≥ 0.5) missense variants, for association with corresponding phenotypes and related Phecodes (Figure 1). Of the 106 genes, 65 genes harbored eligible pLOF or predicted deleterious missense single variants prevalent in PMBB (Table S4). Of these, 18 showed significant evidence of association in the hypothesis-driven single-variant association analyses, namely *AIM1*, *ASPH*, *BRCA1*, *BRCA2*, *CES5A*, *CFTR*, *EPPK1*, *FER1L6*, *FLG2*, *GALC*, *LECT1*, *RTKN2*, *SCNN1D*, *TGM6*, *TTN*, *POLN*, *WDR27*, and *WDR87* (Table 2, Table S5). Importantly, of all 26 gene burdens validated using the two robustness evaluations, three showed evidence of association by both methods (*BRCA1*, *CFTR*, *TGM6*).

### Replication of associations in UK Biobank

We also endeavored to evaluate the robustness of our exome-by-phenome-wide gene burden associations using the UKBB WES dataset, with a specific focus on the 106 pLOF-based gene burden associations with p<E-06 identified in discovery in PMBB. We found that the prevalence of cases for many of the significantly associated phenotypes identified in our discovery phase focusing on pLOF-based gene burden studies in PMBB were significantly lower in UKBB (Table S6), likely due to PMBB being a clinic-based biobank and UKBB being a population-based biobank. For each of the 106 genes, we interrogated 1) gene burdens after collapsing rare pLOF variants, 2) gene burdens after collapsing non-overlapping rare REVEL-predicted deleterious missense variants, and 3) single pLOF or REVEL-predicted deleterious missense variants with MAF > 0.1% on a univariate basis, for hypothesis-driven association studies with their originally associated phenotypes and related Phecodes.

First, 10 of the 106 genes replicated after collapsing rare pLOF variants per gene in UKBB and associating with each gene’s corresponding Phecode and related Phecodes (Table 2, Table S7). Notably, four of these were also significant by the REVEL-informed missense burden or univariate analyses in PMBB (*CYP2D6*, *RGS12*, *TTN*, *WDR27*). Of the other six genes, three were associated with phenotypes with greater prevalence in UKBB versus PMBB. Next, we successfully evaluated the robustness of these associations in UKBB for 14 of the 106 genes by collapsing rare predicted deleterious missense variants per gene (Table 2, Table S8). Seven of these were previously identified in PMBB, and four of seven newly replicated signals were for phenotypes with greater prevalence in UKBB. Finally, 14 of 106 genes were successfully evaluated in UKBB via univariate analyses, of which seven were previously associated in PMBB, and five of seven newly replicated signals were for phenotypes with greater representation in UKBB (Table 2, Table S9). This led to a total of 30 replicated gene-phenotype associations in UKBB, of which 12 had also been validated in PMBB (*ASPH*, *BRCA1*, *CES5A*, *CFTR*, *CYP2D6*, *MYCBP2*, *PPP1R13L*, *RGS12*, *RTKN2*, *TTN*, *WDR27*, *WDR87*). Of note, of the 30 total replicated genes, seven genes showed evidence of association by more than one method in UKBB, of which five also showed evidence of robustness in PMBB (*ASPH*, *CFTR*, *RGS12*, *TTN*, *WDR27*).

### Clinical imaging and laboratory measurements

To build upon our successfully evaluated exome-by-phenome-wide gene-phenotype associations, we took a deeper dive into the cardiovascular imaging and laboratory EHR data in PMBB (Figure 1). First, we analyzed the cardiac structures of carriers for rare pLOF variants in gene burdens associated with cardiomyopathy (*TTN*, *MYBPC3*, *BBS10*) by interrogating their available echocardiography data. As expected based on known biology, carriers of rare pLOF variants in *TTN* had heart morphology consistent with dilated cardiomyopathy when compared to the rest of the PMBB population with echo data available, while carriers of rare pLOFs in *MYBPC3* showed echocardiographic measurements consistent with hypertrophic cardiomyopathy (Table S10). More specifically, *TTN* pLOF carriers had significantly decreased left ventricular outflow tract (LVOT) velocity time integrals and left ventricular ejection fractions, while *MYBPC3* pLOF carriers had significantly increased LVOT velocity time integrals and left ventricular ejection fractions. Despite a lack of power to assess left ventricular function, echocardiography measurements additionally revealed that loss-of-function in *BBS10* is associated with increased LVOT stroke volume, consistent with cardiac hypertrophy, and an increased mitral E/A ratio, indicating supernormal filling (Table S10). Furthermore, interrogation of serum laboratory measurements derived from the EHR showed that *RGS12* is associated with increased serum glucose (ß=65.68, p=1.65E-03).

## Discussion

Here we demonstrate the feasibility and value of aggregating rare pLOF variants into gene burdens on an exome-wide scale for association with a phenome of EHR-derived phenotypes for discovery of novel gene-disease relationships. Using data from 11,451 whole exomes, among the 1518 genes with at least 25 carriers of rare pLOF variants, we identified 106 genes in which a burden of pLOFs was significantly associated with an EHR phenotype. We confirmed the robustness of these associations for 26 genes in PMBB using a separate gene burden of rare (MAF ≤ 0.1%) predicted deleterious missense variants as well as deleterious single variants with MAF > 0.1%, and replicated our findings for 30 genes in UKBB using gene burdens of pLOFs and deleterious variants as well as single variants. In all, a total of 3 positive control gene-disease relationships and 9 novel gene-disease relationships were replicated in both PMBB and UKBB. These results establish that an unbiased gene burden PheWAS in unselected tertiary healthcare biobanks can reveal new gene-disease associations, and given the relatively small size of this experiment suggest that much larger experiments of this type are likely to be highly informative.

Importantly, we identified several positive control gene-disease associations, supporting the validity of our methodology. Our discovery analysis identified a gene burden of rare pLOFs in *CFTR* was significantly associated with cystic fibrosis (CF). CF is a recessive condition caused by biallelic mutations in *CFTR*. We found that the rare pLOF-based gene burden association with CF in PMBB was driven by individuals carrying a second deleterious *CFTR* variant—predominantly deltaF508—that was not included in the pLOF gene burden. This association with CF was not replicated in UKBB due to the extremely low case prevalence of CF (6 cases) compared with PMBB (27 cases). Interestingly, we also found that the *CFTR* pLOF gene burden was associated with bronchiectasis independent of CF and occurred even in individuals without a second *CFTR* mutation, and this finding replicated in UKBB. While a predisposition to bronchiectasis due to haploinsufficiency of *CFTR* has been suggested,^18^ our finding strengthens this observation. *TTN* is a known dilated cardiomyopathy gene and was replicated convincingly in UKBB. *MYBPC3* is a known hypertrophic cardiomyopathy (HCM) gene that showed a convincing association of pLOF burden with HCM in our discovery analysis. Interestingly, it failed to replicate in UKBB where there were only 20 rare pLOF carriers and 12 cases of HCM (*i.e.* Phecodes 425.11 and 425.22), compared with 36 rare pLOF carriers and 185 cases of HCM in PMBB. These results suggest that despite its relatively smaller size compared to UKBB, PMBB has a different—and sicker--population that enables discovery of associations of human diseases driven by rare genetic variants. Finally, *CYP2D6* is a P450 enzyme known to metabolize opiates, and we found that a pLOF gene burden in *CYP2D6* was significantly associated with poisoning by opiates in both PMBB and UKBB.

We identified several novel gene-disease associations in our discovery study that were robust in additional analyses in PMBB as well as replicated in UKBB. Many of these findings have biological plausibility. For example, we found that mutations in *PPP1R13L* are associated with primary open angle glaucoma. *PPP1R13L*, one of the most evolutionarily conserved inhibitors of p53, has been shown to be expressed in the ganglion cell layer of the retina—the most affected tissue in primary open angle glaucoma.^19^ Inhibition of *PPP1R13L* has been shown to exacerbate retinal ganglion cell death following axonal injury^19^ as well as ultraviolet B light-mediated suppression of retinal pigment epithelial cell viability,^20^ further supporting our observation that loss-of-function mutations in *PPP1R13L* increases the risk of primary open angle glaucoma. Additionally, we found that mutations in *RGS12* are associated with severe outcomes of diabetes mellitus. *RGS12* is expressed in islets of Langerhans at both the message and protein level, and has been implicated as a novel cell-cycle related gene in beta cells and is upregulated in replicating beta cells in mice.^21^ While further work must be done to define the functionality of *RGS12* in beta cells, our results in this context strongly suggest a role for loss-of-function mutations in *RGS12* in predisposing to the pathogenesis of diabetes mellitus.

Importantly, gene burden-disease associations with rare phenotypes that were successfully evaluated for robustness in PMBB but not replicated in UKBB may be limited by power in UKBB. For example, a pLOF gene burden in *BBS10* was significantly associated with hypertrophic cardiomyopathy in PMBB and validated by a separate missense gene burden in PMBB; while it was not replicated in UKBB, the prevalence of HCM in UKBB is markedly lower than in PMBB. *BBS10* is one of at least 21 genes implicated in Bardet-Biedl Syndrome, accounting for ∼20% of all cases. Cardiac abnormalities have been reported in Bardet-Biedl Syndrome, including hypertrophy of the interventricular septum.^22^ However, cardiac abnormalities due to haploinsufficiency in *BBS10* have not yet been described.

Given the credibility of these novel associations, we suggest that the lack of replication for associations between rare variants and rare diseases in populations like UKBB that resemble a more normally distributed population implies a greater need for the utilization of healthcare-based academic biobanks that have greater prevalence for rare diseases for discovery of novel gene-disease associations. Finally, while our methodologies for evaluating the robustness of gene-phenotype relationships are valuable when sufficiently powered, we suggest that lack of successful evaluation of robustness within PMBB or in UKBB is likely due to either a lack of power (*i.e.* lack of carriers for other variants in the same genes) or because other variants in those genes may not necessarily be pathogenic or cause comparable effect sizes, and should not discount the initial observations from the pLOF-based exome-by-phenome-wide gene burden studies as false-positive findings.

A significant challenge in rare variant association studies is the difficulty of designing replication studies. While replication of an association using the exact same rare variant in a different population is essentially impossible, we show the value of evaluating the robustness of gene burden associations through complementary, yet non-overlapping genetic association studies from the same cohort, as well as extending to classic replication in independent cohorts. Our approach for evaluating robustness of associations discovered via rare pLOF-based gene burden analyses in PMBB by interrogating other deleterious variants in the same genes (but in different individuals) allowed creating greater confidence in our discovery findings. Given the lack of linkage disequilibrium and mutual carriers between our interrogated rare pLOF variants, rare predicted deleterious missense variants, and low-frequency to common pLOF or predicted deleterious missense variants, we can view each of the three genomic “bins” as nearly-independent events in cases. Because the controls are shared across these analyses, these are considered evaluation of the robustness of the gene-disease relationship rather than true replication. Thus, the probability of identifying association between any two or all three of the bins is highly unlikely to occur due to chance alone. Furthermore, the probability of successfully evaluating the robustness of associations in both PMBB and UKBB is even more unlikely to be due to chance and suggests true biology.

In conclusion, we present an exome-by-phenome-wide association study linking exome sequencing to EHR phenotypes in PMBB, where we focused on rare pLOF variants for discovery of novel gene-phenotype associations, given they are predicted to have the largest effect size compared to other genetic variants. We evaluated the robustness of gene-disease associations by interrogating rare pLOF, rare missense, and single variants in the same gene to address the question of allelic heterogeneity in gene-disease relationships. Our discoveries based in the PMBB dataset supports the use of private, institutional, healthcare-based biobanks for discovery of novel gene-phenotype relationships. Furthermore, our results suggest that this approach applied to even larger cohorts of individuals with whole-exome or whole-genome sequencing data linked to EHR phenotype data will yield many new insights into the relationship of rare genetic variation and human disease phenotypes.

## Acknowledgements

We thank JoEllen Weaver, David Birtwell, Heather Williams, Paul Baumann, and Marjorie Risman.

