## Supplemental Figures and Tables for "Exome-by-phenome-wide rare variant gene burden association with electronic health record phenotypes"

A) B)

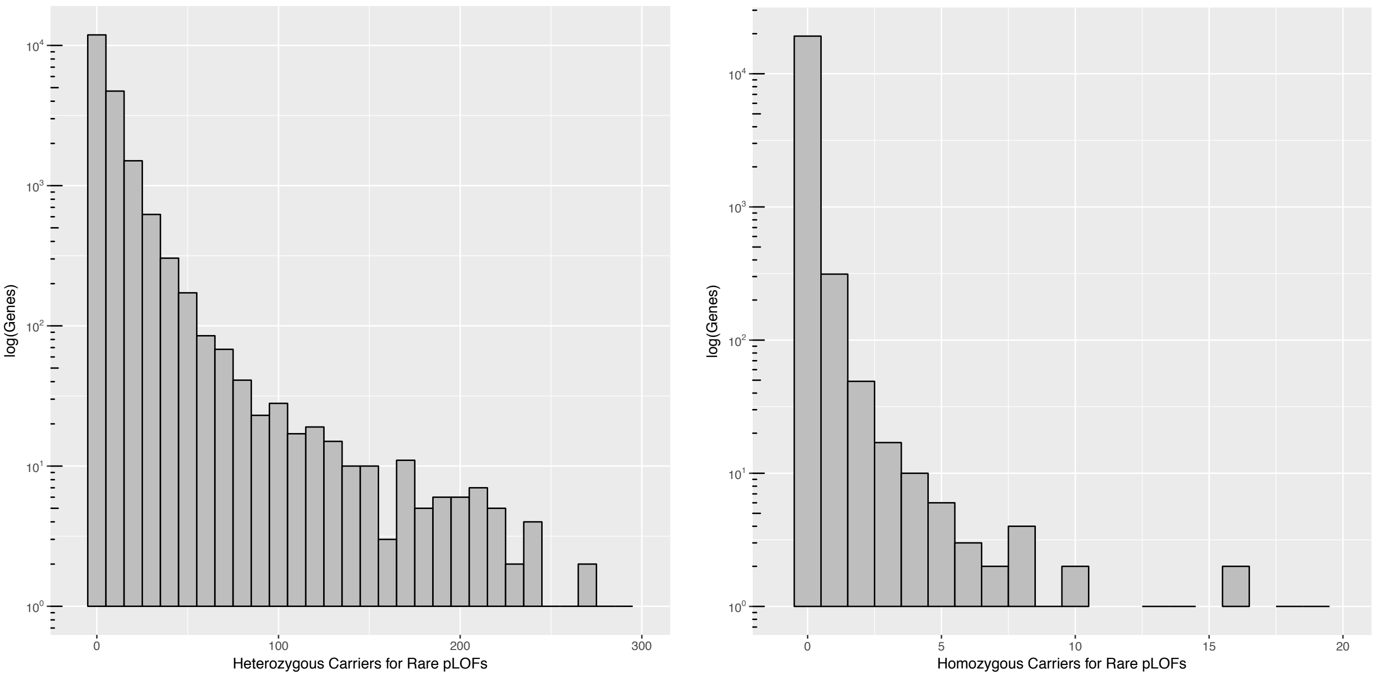


**Figure S1. Distribution of number of carriers for predicted loss-of-function (pLOF) variants per gene in Penn Medicine Biobank (PMBB)**

A) Histogram plot for the distribution of number of heterozygous carriers for pLOF variants per gene in PMBB's exome sequenced cohort. The x-axis represents number of heterozygous pLOF carriers per gene in bin widths of 10, and the log-scaled y-axis represents the number of genes with the x-axis-specified number of heterozygous carriers. B) Histogram plot for the distribution of number of homozygous carriers for pLOF variants per gene in PMBB's exome sequenced cohort. The x-axis represents number of homozygous pLOF carriers per gene in bin widths of one, and the log-scaled y-axis represents the number of genes with the x-axis-specified number of homozygous carriers.

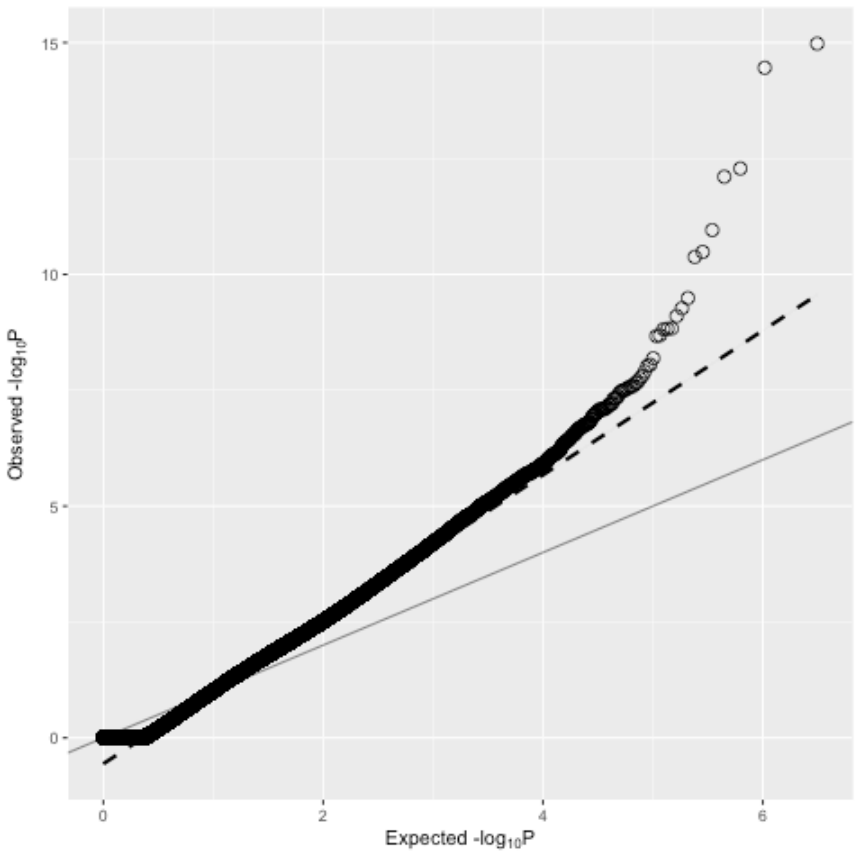


**Figure S2. Quantile-quantile plot of gene burden testing results from discovery phase of exome-by-phenome-wide association studies in Penn Medicine Biobank.**

Quantile-quantile plot of p values from exome-by-phenome-wide association studies using gene burdens collapsing rare (MAF ≤ 0.1%) predicted loss-of-function (pLOF) variants per gene in Penn Medicine Biobank (PMBB). The x-axis represents the expected $-\log_{10}$(p value) under the uniform distribution of p values. The y axis represents the observed $-\log_{10}$(p value) from the discovery phase of the exome-by-phenome-wide gene burden association studies collapsing rare pLOF variants in PMBB. Each point represents an association between one of 1518 gene burdens and one of 1000 Phecodes. The solid line shows the relationship between the expected and observed p values under the uniform p value distribution. The dashed line represents the observed fit line between the 50$^{th}$ and 95$^{th}$ percentile of gene burden associations, and the slope of this line is $\lambda_{\Delta 95} = 1.558$.

### Table S1

| Gene | Phecode | Phecode Description | Logistic ß | Logistic p | Firth's ß | Firth's p |
|---|---|---|---|---|---|---|
| *ABCA10* | 225 | Benign neoplasm of brain and other parts of nervous system | 3.546 | 1.11E-07 | 3.575 | 2.81E-10 |
| | 225.1 | Benign neoplasm of brain, cranial nerves, meninges | 3.599 | 7.26E-08 | 3.669 | 7.56E-11 |
| | 514.1 | Abnormal results of function study of pulmonary system | 3.604 | 1.54E-07 | 3.375 | 9.34E-09 |
| *ADM* | 444 | Arterial embolism and thrombosis | 1.181 | 7.87E-07 | 1.134 | 2.92E-07 |
| *AIM1* | 741 | Symptoms and disorders of the joints | 2.424 | 3.05E-07 | 2.492 | 5.20E-08 |
| *AK9* | 274.11 | Gouty arthropathy | 3.096 | 9.38E-09 | 2.871 | 9.01E-09 |
| *AKR1C3* | 962.1 | Adrenal cortical steroids causing adverse effects in therapeutic use | 3.107 | 6.28E-07 | 2.826 | 7.62E-07 |
| *ALPK1* | 735.21 | Hammer toe (acquired) | 3.107 | 7.06E-07 | 2.985 | 1.98E-07 |
| *AMHR2* | 296.1 | Bipolar | 3.839 | 6.84E-07 | 2.614 | 3.69E-06 |
| *ANKRD35* | 521.1 | Dental caries | 3.123 | 8.84E-07 | 2.738 | 1.33E-07 |
| *ASPH* | 681.3 | Cellulitis and abscess of arm/hand | 4.305 | 5.39E-10 | 4.065 | 1.81E-11 |
| *ATP7B* | 251 | Other disorders of pancreatic internal secretion | 3.401 | 3.30E-07 | 3.449 | 1.49E-08 |
| *BAIAP2L2* | 334 | Degenerative disease of the spinal cord | 2.549 | 8.38E-08 | 2.418 | 1.47E-08 |
| *BAIAP3* | 246.7 | Abnormal results of function study of thyroid | 2.837 | 7.15E-07 | 2.786 | 1.25E-07 |
| *BBS10* | 425.11 | Hypertrophic obstructive cardiomyopathy | 2.927 | 2.89E-08 | 2.904 | 4.25E-09 |
| *BRCA1* | 433.6 | Acute, but ill-defined cerebrovascular disease | 3.718 | 3.21E-08 | 3.482 | 1.31E-08 |
| *BRCA2* | 174 | Breast cancer | 2.228 | 1.72E-07 | 2.200 | 1.76E-07 |
| | 174.1 | Breast cancer [female] | 2.228 | 1.88E-07 | 2.198 | 1.96E-07 |
| | 174.11 | Malignant neoplasm of female breast | 2.261 | 1.36E-07 | 2.230 | 1.44E-07 |
| | 368.2 | Diplopia and disorders of binocular vision | 2.802 | 7.99E-07 | 2.845 | 1.64E-08 |
| *BRSK1* | 345.1 | Epilepsy | 3.289 | 3.10E-07 | 2.661 | 5.43E-07 |
| | 345.12 | Partial epilepsy | 3.767 | 3.01E-08 | 2.961 | 6.54E-08 |
| *BTN2A1* | 729 | Other disorders of soft tissues | 3.735 | 2.77E-07 | 3.767 | 1.24E-08 |
| *C14ORF183* | 260.6 | Anorexia | 4.031 | 9.42E-08 | 3.910 | 3.03E-09 |
| *CCDC144NL* | 500.2 | Pneumoconiosis | 4.956 | 9.45E-07 | 4.793 | 5.69E-09 |
| *CCDC18* | 359 | Muscular dystrophies and other myopathies | 3.990 | 9.46E-07 | 3.671 | 6.20E-08 |
| | 359.2 | Myopathy | 4.225 | 3.89E-07 | 3.928 | 1.42E-08 |
| *CCDC60* | 198.4 | Secondary malignant neoplasm of liver | 3.253 | 8.66E-07 | 3.356 | 9.64E-09 |
| *CCDC74A* | 198.1 | Secondary malignancy of lymph nodes | 3.452 | 2.05E-07 | 3.359 | 9.30E-09 |
| *CENPF* | 382 | Otalgia | 3.225 | 9.20E-07 | 3.258 | 3.32E-08 |
| *CES5A* | 286.9 | Abnormal coagulation profile | 3.306 | 8.10E-08 | 3.204 | 4.52E-08 |
| *CFTR* | 480.12 | Pseudomonal pneumonia | 2.985 | 2.27E-07 | 2.967 | 4.92E-09 |
| | 496.3 | Bronchiectasis | 3.044 | 4.21E-11 | 2.939 | 5.26E-12 |
| | 499 | Cystic fibrosis | 4.447 | 1.05E-15 | 4.280 | 2.26E-18 |
| *CILP* | 447.7 | Aortic ectasia | 2.946 | 4.29E-08 | 3.015 | 5.54E-09 |
| *CPT1C* | 447 | Other disorders of arteries and arterioles | 2.221 | 2.01E-07 | 2.065 | 5.10E-07 |
| *CRIPAK* | 571.6 | Primary biliary cirrhosis | 4.465 | 7.69E-07 | 3.945 | 2.19E-08 |

| | | | | | | |
|---|---|---|---|---|---|---|
| *CTC1* | 526.4 | Temporomandibular joint disorders | 4.415 | 3.76E-07 | 4.466 | 1.24E-09 |
| *CYP2D6* | 965.1 | Opiates and related narcotics causing adverse effects in therapeutic use | 2.743 | 1.50E-09 | 2.773 | 8.34E-11 |
| *DDX60L* | 687.4 | Disturbance of skin sensation | 2.354 | 9.31E-07 | 2.002 | 1.13E-05 |
| *DNAH6* | 350.3 | Lack of coordination | 3.646 | 7.93E-10 | 3.623 | 8.59E-14 |
| *DNHD1* | 733.4 | Aseptic necrosis of bone | 2.879 | 2.67E-07 | 2.796 | 6.80E-09 |
| *DOCK6* | 31 | Diseases due to other mycobacteria | 4.309 | 2.89E-07 | 4.028 | 2.14E-09 |
| *EFCAB5* | 281 | Other deficiency anemia | 3.359 | 8.31E-08 | 3.314 | 1.97E-08 |
| | 618.1 | Prolapse of vaginal walls | 4.288 | 3.19E-08 | 4.230 | 6.21E-06 |
| *EFCAB6* | 271 | Disorders of carbohydrate transport and metabolism | 4.057 | 7.72E-07 | 4.083 | 1.32E-04 |
| *EPPK1* | 451.2 | Phlebitis and thrombophlebitis of lower extremities | 2.698 | 9.19E-08 | 2.410 | 6.50E-08 |
| *EXTL2* | 289 | Other diseases of blood and blood-forming organs | 5.327 | 6.15E-08 | 5.257 | 1.19E-05 |
| *FARSA* | 496.2 | Chronic bronchitis | 3.221 | 7.38E-07 | 2.928 | 1.30E-06 |
| | 800 | Fracture of lower limb | 3.344 | 8.03E-07 | 3.128 | 3.10E-07 |
| *FER1L6* | 772.1 | Muscular wasting and disuse atrophy | 3.197 | 7.18E-07 | 3.110 | 1.63E-08 |
| *FLG2* | 741.2 | Stiffness of joint | 3.173 | 1.76E-07 | 3.080 | 6.95E-09 |
| *FNDC4* | 698 | Pruritus and related conditions | 2.728 | 4.73E-08 | 2.607 | 1.01E-08 |
| *FREM1* | 560.4 | Other intestinal obstruction | 3.563 | 2.12E-07 | 3.312 | 9.45E-08 |
| *GALC* | 596 | Other disorders of bladder | 2.559 | 3.48E-07 | 2.573 | 1.52E-07 |
| *GDAP2* | 425.12 | Other hypertrophic cardiomyopathy | 3.817 | 1.46E-09 | 3.804 | 1.61E-06 |
| *GHDC* | 742 | Derangement of joint, non-traumatic | 4.347 | 2.43E-07 | 3.724 | 1.23E-08 |
| | 742.9 | Other derangement of joint | 4.536 | 1.35E-07 | 3.915 | 5.52E-09 |
| *GTF2I* | 574.3 | Cholecystitis without cholelithiasis | 3.875 | 1.24E-08 | 3.687 | 7.93E-10 |
| *HEATR4* | 964 | Poisoning by agents primarily affecting blood constituents | 3.349 | 9.27E-07 | 3.263 | 9.35E-08 |
| | 964.1 | Anticoagulants causing adverse effects | 3.431 | 5.82E-07 | 3.329 | 5.81E-08 |
| *HPS1* | 604 | Disorders of penis | 3.908 | 2.73E-07 | 3.565 | 6.47E-08 |
| *IL20* | 442.3 | Aneurysm of artery of lower extremity | 3.699 | 4.08E-07 | 3.686 | 2.16E-08 |
| *IPP* | 870 | Open wounds of head; neck; and trunk | 3.305 | 4.65E-08 | 3.046 | 3.17E-08 |
| *IQCE* | 528 | Diseases of the oral soft tissues, excluding lesions specific for gingiva and tongue | 2.787 | 3.80E-07 | 2.701 | 1.02E-07 |
| *KCNH2* | 8.6 | Viral Enteritis | 1.964 | 1.07E-07 | 1.756 | 1.54E-08 |
| *KIAA0319* | 938.2 | Chronic dermatitis due to solar radiation | 3.499 | 7.90E-07 | 3.385 | 4.52E-08 |
| *KIAA1755* | 990 | Effects radiation NOS | 6.163 | 5.19E-07 | 3.893 | 1.47E-05 |
| *LECT1* | 555.2 | Ulcerative colitis | 3.297 | 7.11E-07 | 3.423 | 2.43E-08 |
| *LIPE* | 870.3 | Other open wound of head and face | 4.163 | 2.83E-07 | 4.062 | 3.31E-09 |
| *LRRCC1* | 571.81 | Portal hypertension | 3.684 | 4.63E-07 | 3.555 | 6.83E-08 |
| *MAD1L1* | 681.6 | Cellulitis and abscess of foot, toe | 4.339 | 3.94E-07 | 3.427 | 2.03E-07 |
| *MICALL2* | 681.6 | Cellulitis and abscess of foot, toe | 4.351 | 7.75E-07 | 4.378 | 4.42E-09 |
| *MUC17* | 450 | Noninfectious disorders of lymphatic channels | 1.704 | 2.59E-08 | 1.706 | 4.76E-09 |
| *MYBPC3* | 425.11 | Hypertrophic obstructive cardiomyopathy | 3.675 | 5.20E-13 | 3.632 | 4.88E-13 |
| | 425.12 | Other hypertrophic cardiomyopathy | 4.265 | 3.49E-15 | 4.237 | 5.54E-16 |
| | 574.3 | Cholecystitis without cholelithiasis | 3.592 | 8.23E-08 | 3.421 | 3.43E-09 |

| | | | | | | |
|---|---|---|---|---|---|---|
| *MYCBP2* | 772.2 | Spasm of muscle | 3.411 | 2.08E-07 | 3.459 | 4.96E-05 |
| *MYO1A* | 689 | Disorder of skin and subcutaneous tissue NOS | 2.772 | 2.68E-08 | 2.774 | 2.30E-09 |
| *OR11H1* | 145 | Cancer of mouth | 3.578 | 1.67E-07 | 3.581 | 2.77E-09 |
| | 145.2 | Cancer of tongue | 4.335 | 2.07E-09 | 4.347 | 6.89E-12 |
| *OR5H6* | 771.2 | Cramp of limb | 1.710 | 9.59E-07 | 1.762 | 7.65E-08 |
| *PER3* | 201 | Hodgkin's disease | 3.804 | 4.80E-08 | 3.777 | 1.81E-09 |
| *PIWIL3* | 500.2 | Pneumoconiosis | 5.025 | 1.04E-07 | 4.880 | 2.01E-10 |
| | 574.2 | Calculus of bile duct | 3.808 | 2.13E-08 | 3.608 | 2.10E-09 |
| *PLSCR5* | 711 | Arthropathy associated with infections | 4.223 | 4.38E-07 | 4.073 | 2.29E-09 |
| *POLN* | 289 | Other diseases of blood and blood-forming organs | 3.859 | 2.57E-08 | 3.581 | 2.87E-09 |
| | 572 | Ascites (non malignant) | 2.593 | 1.55E-07 | 2.604 | 3.79E-08 |
| *PPM1D* | 300.12 | Agorophobia, social phobia, and panic disorder | 3.657 | 6.49E-07 | 3.679 | 7.24E-08 |
| *PPP1R13L* | 365.11 | Primary open angle glaucoma | 3.117 | 7.29E-07 | 3.187 | 3.68E-08 |
| *PRDM2* | 361 | Retinal detachments and defects | 2.448 | 6.28E-07 | 2.459 | 1.22E-08 |
| *PSG11* | 514.1 | Abnormal results of function study of pulmonary system | 5.178 | 4.90E-07 | 4.451 | 1.50E-08 |
| *PSG8* | 454.11 | Varicose veins of lower extremity, symptomtic | 3.461 | 8.20E-08 | 3.247 | 6.28E-08 |
| *RGS12* | 250.12 | Type 1 diabetes with renal manifestations | 3.874 | 6.48E-08 | 3.642 | 6.84E-09 |
| | 250.13 | Type 1 diabetes with ophthalmic manifestations | 4.042 | 3.36E-08 | 3.794 | 1.67E-09 |
| *RNPEP* | 378 | Strabismus and other disorders of binocular eye movements | 3.403 | 8.10E-07 | 3.458 | 2.02E-08 |
| *RSU1* | 558 | Noninfectious gastroenteritis | 1.718 | 7.51E-08 | 1.753 | 1.34E-08 |
| | 715 | Other inflammatory spondylopathies | 2.038 | 6.82E-07 | 2.061 | 2.57E-08 |
| *RTKN2* | 458.1 | Orthostatic hypotension | 4.436 | 7.24E-07 | 3.730 | 9.54E-07 |
| *SCNN1D* | 426.8 | Other cardiac conduction disorders | 4.189 | 4.52E-07 | 3.601 | 8.82E-07 |
| *SCRN3* | 446 | Polyarteritis nodosa and allied conditions | 3.313 | 6.63E-07 | 3.313 | 4.42E-08 |
| *SEC14L3* | 536.3 | Gastroparesis | 3.268 | 6.21E-08 | 3.251 | 8.80E-09 |
| *SH3D21* | 733.4 | Aseptic necrosis of bone | 4.043 | 2.02E-07 | 3.956 | 1.30E-08 |
| | 735.3 | Hallux valgus (Bunion) | 3.723 | 2.32E-07 | 3.767 | 9.96E-09 |
| *SIGLECL1* | 274 | Gout and other crystal arthropathies | 1.855 | 4.57E-07 | 1.866 | 4.98E-06 |
| | 274.1 | Gout | 1.863 | 4.21E-07 | 1.874 | 4.66E-06 |
| *SLC12A3* | 442.3 | Aneurysm of artery of lower extremity | 4.513 | 3.28E-11 | 4.251 | 1.91E-12 |
| *SLC22A2* | 508 | Pulmonary collapse; interstitial and compensatory emphysema | 3.377 | 8.06E-07 | 3.206 | 1.32E-07 |
| *SLC2A8* | 726.2 | Synoviopathy | 3.586 | 1.83E-07 | 3.401 | 1.00E-08 |
| *SPG7* | 130 | Spirochetal infection | 3.504 | 2.14E-09 | 3.389 | 6.77E-11 |
| | 130.1 | Lyme disease | 3.541 | 1.53E-09 | 3.432 | 2.96E-11 |
| *STOX2* | 365.1 | Open-angle glaucoma | 1.947 | 3.50E-07 | 1.916 | 2.62E-07 |
| *SYT15* | 430 | Intracranial hemorrhage | 2.626 | 1.81E-07 | 2.587 | 2.10E-08 |
| *TGM6* | 214 | Lipoma | 2.571 | 2.77E-07 | 2.401 | 1.23E-07 |
| *TMCO1* | 735.3 | Hallux valgus (Bunion) | 3.746 | 1.60E-07 | 3.575 | 1.51E-08 |
| | 830 | Dislocation | 3.118 | 1.16E-07 | 2.970 | 5.65E-08 |
| *TRDN* | 735.2 | Acquired toe deformities | 3.311 | 3.90E-07 | 2.964 | 1.51E-07 |

| | | | | | | |
|---|---|---|---|---|---|---|
| *TTC21B* | 377 | Disorders of optic nerve and visual pathways | 4.051 | 8.50E-07 | 3.179 | 9.99E-08 |
| *TTN* | 425 | Cardiomyopathy | 0.876 | 7.83E-13 | 0.876 | 7.84E-13 |
| | 425.1 | Primary/intrinsic cardiomyopathies | 0.874 | 1.11E-11 | 0.875 | 1.02E-11 |
| | 425.2 | Secondary/extrinsic cardiomyopathies | 0.899 | 9.28E-07 | 0.910 | 5.10E-07 |
| | 426.9 | Cardiac pacemaker/device in situ | 0.846 | 3.69E-07 | 0.839 | 4.44E-07 |
| | 426.92 | Cardiac defibrillator in situ | 0.989 | 1.77E-08 | 0.982 | 2.09E-08 |
| | 427.1 | Paroxysmal tachycardia, unspecified | 0.942 | 6.45E-09 | 0.933 | 8.76E-09 |
| | 427.12 | Paroxysmal ventricular tachycardia | 0.956 | 2.18E-08 | 0.946 | 2.88E-08 |
| | 427.6 | Premature beats | 1.113 | 8.90E-09 | 1.109 | 8.43E-09 |
| *UBALD2* | 705.8 | Hyperhidrosis | 2.242 | 4.15E-07 | 2.277 | 4.24E-08 |
| *WDR27* | 695.7 | Prurigo and Lichen | 4.168 | 6.31E-07 | 3.897 | 2.65E-08 |
| *WDR87* | 550.5 | Ventral hernia | 2.122 | 1.70E-07 | 1.997 | 9.58E-08 |
| *ZC3H3* | 573 | Other disorders of liver | 2.414 | 1.56E-08 | 2.408 | 2.67E-09 |
| *ZNF175* | 389.4 | Tinnitus | 3.347 | 3.24E-10 | 3.380 | 3.02E-11 |
| *ZNF266* | 356 | Hereditary and idiopathic peripheral neuropathy | 3.385 | 3.63E-08 | 3.264 | 1.12E-08 |
| *ZNF334* | 593.2 | Microscopic hematuria | 3.078 | 1.69E-07 | 2.768 | 1.24E-07 |
| | 772.3 | Muscle weakness | 3.365 | 3.22E-07 | 3.421 | 5.72E-05 |
| *ZNF354C* | 565 | Anal and rectal conditions | 3.100 | 1.03E-07 | 3.089 | 2.00E-08 |
| *ZNF582* | 689 | Disorder of skin and subcutaneous tissue NOS | 3.114 | 8.00E-08 | 3.223 | 1.92E-09 |
| *ZNF587* | 145 | Cancer of mouth | 3.517 | 2.17E-07 | 3.576 | 4.59E-09 |
| | 145.2 | Cancer of tongue | 3.849 | 6.49E-08 | 4.017 | 3.21E-10 |
| *ZNF837* | 695.7 | Prurigo and Lichen | 3.052 | 6.13E-07 | 2.175 | 1.24E-06 |

**Table S1. Summary statistics for 106 gene burdens with associations with p<E-06 from exome-by-phenome-wide association studies in Penn Medicine Biobank.**

List of 139 associations across 106 unique gene burdens with p<E-06 from exome-by-phenome-wide association studies in Penn Medicine Biobank based on an exact logistic regression model adjusted for age, $age^2$, gender, and the first ten principal components of ancestry. Reported are the betas and p-values resulting from meta-analysis of the summary statistics obtained from running the logistic regression model separately in individuals of European and African ancestry. Also reported are the betas and p-values resulting from meta-analysis of European- and African-specific summary statistics based on Firth's penalized likelihood model, also adjusted for age, $age^2$, gender, and the first ten principal components of ancestry.

**Table S2**

| Gene | Homozygous N | Heterozygous N |
|---|---|---|
| *ABCA10* | 0 | 27 |
| *ADM* | 0 | 0 |
| *AIM1* | 0 | 62 |
| *AK9* | 0 | 34 |
| *AKR1C3* | 0 | 8 |
| *ALPK1* | 0 | 6 |
| *AMHR2* | 0 | 11 |
| *ANKRD35* | 0 | 29 |
| *ASPH* | 0 | 30 |
| *ATP7B* | 0 | 257 |
| *BAIAP2L2* | 0 | 12 |
| *BAIAP3* | 0 | 86 |
| *BBS10** | 0 | 23 |
| *BRCA1** | 0 | 153 |
| *BRCA2* | 0 | 112 |
| *BRSK1* | 0 | 0 |
| *BTN2A1* | 0 | 4 |
| *C14ORF183* | 0 | 0 |
| *CCDC144NL* | 0 | 0 |
| *CCDC18* | 0 | 0 |
| *CCDC60* | 0 | 0 |
| *CCDC74A* | 0 | 1 |
| *CENPF* | 0 | 5 |
| *CES5A* | 0 | 33 |
| *CFTR** | 1 | 563 |
| *CILP** | 0 | 33 |
| *CPT1C* | 0 | 59 |
| *CRIPAK* | 0 | 0 |
| *CTC1* | 2 | 72 |
| *CYP2D6** | 0 | 40 |
| *DDX60L* | 0 | 2 |
| *DNAH6* | 0 | 87 |
| *DNHD1* | 0 | 39 |
| *DOCK6* | 0 | 84 |
| *EFCAB5* | 0 | 1 |
| *EFCAB6* | 0 | 9 |
| *EPPK1* | 1 | 210 |
| *EXTL2* | 0 | 59 |
| *FARSA* | 0 | 38 |
| *FER1L6* | 0 | 196 |
| *FLG2* | 0 | 0 |

| | | |
|---|---|---|
| *FNDC4* | 0 | 3 |
| *FREM1* | 0 | 50 |
| *GALC* | 0 | 106 |
| *GDAP2* | 0 | 5 |
| *GHDC* | 0 | 13 |
| *GTF2I* | 0 | 3 |
| *HEATR4* | 0 | 0 |
| *HPS1* | 0 | 13 |
| *IL20* | 0 | 5 |
| *IPP* | 0 | 27 |
| *IQCE* | 0 | 5 |
| *KCNH2** | 1 | 314 |
| *KIAA0319* | 0 | 13 |
| *KIAA1755* | 0 | 2 |
| *LECT1* | 0 | 11 |
| *LIPE* | 0 | 38 |
| *LRRCC1* | 0 | 4 |
| *MAD1L1* | 0 | 6 |
| *MICALL2* | 1 | 39 |
| *MUC17* | 0 | 0 |
| *MYBPC3** | 0 | 125 |
| *MYCBP2** | 0 | 74 |
| *MYO1A* | 0 | 140 |
| *OR11H1* | 0 | 0 |
| *OR5H6* | 0 | 0 |
| *PER3* | 0 | 3 |
| *PIWIL3* | 0 | 0 |
| *PLSCR5* | 0 | 11 |
| *POLN* | 0 | 53 |
| *PPM1D* | 0 | 0 |
| *PPP1R13L** | 0 | 10 |
| *PRDM2* | 0 | 9 |
| *PSG11* | 0 | 0 |
| *PSG8* | 0 | 1 |
| *RGS12** | 0 | 32 |
| *RNPEP* | 0 | 18 |
| *RSU1* | 0 | 3 |
| *RTKN2* | 0 | 7 |
| *SCNN1D* | 0 | 4 |
| *SCRN3** | 0 | 18 |
| *SEC14L3* | 1 | 20 |
| *SH3D21* | 0 | 3 |
| *SIGLECL1* | 0 | 2 |
| *SLC12A3* | 0 | 180 |
| *SLC22A2* | 0 | 44 |

| | | |
|---|---|---|
| *SLC2A8* | 0 | 90 |
| *SPG7* | 1 | 122 |
| *STOX2* | 0 | 25 |
| *SYT15* | 0 | 11 |
| *TGM6** | 0 | 117 |
| *TMCO1* | 0 | 3 |
| *TRDN* | 0 | 2 |
| *TTC21B* | 0 | 93 |
| *TTN* | 4 | 1264 |
| *UBALD2* | 0 | 0 |
| *WDR27* | 0 | 20 |
| *WDR87* | 0 | 5 |
| *ZC3H3* | 0 | 3 |
| *ZNF175* | 0 | 2 |
| *ZNF266* | 0 | 0 |
| *ZNF334* | 0 | 1 |
| *ZNF354C* | 0 | 3 |
| *ZNF582* | 0 | 0 |
| *ZNF587* | 0 | 0 |
| *ZNF837* | 0 | 0 |

**Table S2. Number of carriers for rare predicted deleterious missense variants among 106 genes of interest.**

List of 106 genes with associations p<E-06 from exome-by-phenome-wide association studies using pLOF-based gene burdens. Each gene is labeled with the number of homozygous and heterozygous carriers for rare (minor allele frequency ≤ 0.1% per gnomAD among Europeans and Africans) missense variants with REVEL scores of at least 0.5 that were prevalent in PMBB. Note: starred genes (*) represent those for which their REVEL-informed missense-based gene burden studies successfully replicated associations identified from the pLOF-based gene burden studies.

### Table S3

| | pLOF-Based Gene Burden | | | | | REVEL-Informed Missense-Based Gene Burden | | | | |
|---|---|---|---|---|---|---|---|---|---|---|
| Gene | Phecode | Description | OR | P | N | Phecode | Description | OR | P | N |
| *BBS10* | 425.11 | Hypertrophic obstructive cardiomyopathy | 18.66 | 2.89E-08 | 39 het | 425.11 | Hypertrophic obstructive cardiomyopathy | 8.83 | 4.12E-02 | 23 het |
| | | | | | | 425.12 | Other hypertrophic cardiomyopathy | 10.25 | 3.18E-02 | |
| *BRCA1* | 433.6 | Acute, but ill-defined cerebrovascular disease | 41.19 | 3.21E-08 | 25 het | 433.1 | Occlusion and stenosis of precerebral arteries | 1.82 | 3.60E-02 | 149 het, 2 double het |
| *CFTR* | 499 | Cystic fibrosis | 85.39 | 1.05E-15 | 84 het | 499 | Cystic fibrosis | 13.50 | 6.24E-11 | 1 hom, 538 het, 11 double het, 1 triple het |
| | 496.3 | Bronchiectasis | 20.99 | 4.21E-11 | | 496.3 | Bronchiectasis | 2.71 | 4.33E-03 | |
| | 480.12 | Pseudomonal pneumonia | 19.79 | 2.27E-07 | | 480.12 | Pseudomonal pneumonia | 4.03 | 1.30E-04 | |
| *CILP* | 447.7 | Aortic ectasia | 19.04 | 4.29E-08 | 32 het | 447.7 | Aortic ectasia | 9.70 | 5.05E-04 | 33 het |
| *CYP2D6* | 965.1 | Opiates and related narcotics causing adverse effects in therapeutic use | 15.54 | 1.50E-09 | 1 hom, 100 het | 965.1 | Opiates and related narcotics causing adverse effects in therapeutic use | 11.71 | 2.05E-02 | 40 het |
| *KCNH2* | 8.6 | Viral Enteritis | 7.13 | 1.07E-07 | 181 het, 40 double het | 8.52 | Intestinal infection due to C. difficile | 2.30 | 2.45E-02 | 1 hom, 290 het, 12 double het |
| *MYBPC3* | 425.12 | Other hypertrophic cardiomyopathy | 71.14 | 3.49E-15 | 36 het | 425.12 | Other hypertrophic cardiomyopathy | 4.49 | 1.49E-02 | 123 single het, 1 double het |
| | 425.11 | Hypertrophic obstructive cardiomyopathy | 39.45 | 5.20E-13 | | | | | | |
| *MYCBP2* | 772.2 | Spasm of muscle | 30.28 | 2.08E-07 | 33 het | 772.2 | Spasm of muscle | 16.94 | 2.22E-05 | 74 het |
| *PPP1R13L* | 365.11 | Primary open angle glaucoma | 22.58 | 7.29E-07 | 41 het | 365.11 | Primary open angle glaucoma | 38.64 | 2.07E-02 | 10 het |
| *RGS12* | 250.13 | Type 1 diabetes with ophthalmic manifestations | 56.94 | 3.36E-08 | 41 het | 250.23 | Type 2 diabetes with ophthalmic manifestations | 5.99 | 5.52E-03 | 32 het |
| | 250.12 | Type 1 diabetes with renal manifestations | 48.14 | 6.48E-08 | | 250.7 | Diabetic retinopathy | 6.25 | 1.92E-02 | |
| *SCRN3* | 446 | Polyarteritis nodosa and allied conditions | 27.48 | 6.63E-07 | 27 het | 446 | Polyarteritis nodosa and allied conditions | 30.39 | 3.49E-03 | 18 het |
| *TGM6* | 214 | Lipoma | 13.08 | 2.77E-07 | 17 het, 5 double het | 214 | Lipoma | 4.20 | 4.51E-02 | 115 single het, 1 double het |

**Table S3: Evaluation of robustness via REVEL-informed missense-based gene burdens within Penn Medicine Biobank.**

List of significantly replicated associations via gene burdens collapsing rare (MAF ≤ 0.1%) predicted deleterious missense (REVEL ≥ 0.5) variants in Penn Medicine Biobank (PMBB) among 106 genes for which predicted loss-of-function (pLOF)-based gene burdens had p<E-06 from exome-by-phenome-wide association studies in PMBB. Summary statistics are based on an exact logistic regression model adjusted for age, $age^2$, gender, and the first ten principal components of ancestry. Reported are the odds ratios and p-values resulting from meta-analysis of the summary statistics obtained from running the logistic regression model separately in individuals of European and African ancestry, and the number of carriers for each gene burden.

### Table S4

| Gene | Chr | Position | Ref | Alt | Variant Type | gnomAD AFR MAF | gnomAD NFE MAF | REVEL | Hom | Het |
|---|---|---|---|---|---|---|---|---|---|---|
| *ABCA10** | 17 | 67149973 | G | A | stopgain | 0.0757 | 0.0504 | NA | 57 | 1296 |
| | 17 | 67150465 | - | ACCTG GAA | frameshift insertion | 0.3172 | 9.00E-04 | NA | 217 | 1018 |
| | 17 | 67170519 | C | T | nonsynonymous SNV | 0.0035 | 1.88E-05 | 0.57 | 0 | 13 |
| | 17 | 67178422 | A | T | nonsynonymous SNV | 0.0023 | 2.06E-05 | 0.503 | 0 | 9 |
| | 17 | 67189302 | G | A | stopgain | 0.0434 | 2.00E-04 | NA | 3 | 180 |
| | 17 | 67190568 | T | C | nonsynonymous SNV | 0.0096 | 0 | 0.655 | 0 | 37 |
| | 17 | 67190619 | A | G | nonsynonymous SNV | 0.0063 | 9.11E-06 | 0.762 | 0 | 26 |
| | 17 | 67211985 | G | T | nonsynonymous SNV | 0.019 | 2.00E-04 | 0.511 | 1 | 81 |
| | 17 | 67212031 | A | T | stopgain | 0.1983 | 0.0012 | NA | 75 | 724 |
| | 17 | 67215702 | G | A | stopgain | 0.0125 | 9.16E-05 | NA | 0 | 59 |
| | 17 | 67218672 | A | G | splicing | 0.0018 | 0 | NA | 0 | 6 |
| *AIM1** | 6 | 106991361 | T | C | nonsynonymous SNV | 0.011 | 0.0598 | 0.729 | 37 | 1100 |
| | 6 | 106992472 | T | C | nonsynonymous SNV | 0.0173 | 0 | 0.816 | 1 | 67 |
| | 6 | 106992477 | G | A | nonsynonymous SNV | 0.0011 | 2.69E-05 | 0.909 | 0 | 8 |
| | 6 | 107003686 | G | A | nonsynonymous SNV | 0.0016 | 0.0016 | 0.566 | 0 | 42 |
| *AK9* | 6 | 109894799 | - | A | splicing | 0.0227 | 7.70E-05 | NA | 2 | 108 |
| *AKR1C3* | 10 | 5141639 | C | T | stopgain | 0.0058 | 2.00E-04 | NA | 0 | 23 |
| | 10 | 5141642 | G | A | splicing | 0.0049 | 8.96E-06 | NA | 0 | 27 |
| | 10 | 5144336 | - | A | frameshift insertion | 0.0028 | 0 | NA | 0 | 17 |
| *ALPK1* | 4 | 113352488 | G | A | stopgain | 0.0087 | 1.79E-05 | NA | 0 | 35 |
| *ANKRD35* | 1 | 145561822 | C | T | stopgain | 0.0013 | 3.59E-05 | NA | 0 | 11 |
| *ASPH** | 8 | 62550924 | - | A | splicing | 0.2945 | 0.3109 | NA | 87 | 6623 |
| | 8 | 62550924 | - | AA | splicing | 0.0164 | 0.0103 | NA | 0 | 481 |
| *ATP7B* | 13 | 52509155 | G | A | nonsynonymous SNV | 4.00E-04 | 0.0018 | 0.879 | 0 | 33 |
| | 13 | 52518281 | G | T | nonsynonymous SNV | 0 | 0.0013 | 0.909 | 0 | 35 |
| | 13 | 52520508 | G | A | nonsynonymous SNV | 5.00E-04 | 0.0024 | 0.927 | 0 | 33 |
| | 13 | 52524268 | C | T | nonsynonymous SNV | 2.00E-04 | 0.0012 | 0.876 | 0 | 24 |
| | 13 | 52534410 | C | T | nonsynonymous SNV | 5.00E-04 | 0.0017 | 0.655 | 1 | 59 |
| | 13 | 52542680 | A | G | nonsynonymous SNV | 7.00E-04 | 0.0043 | 0.555 | 0 | 74 |
| | 13 | 52542732 | C | T | nonsynonymous SNV | 1.00E-04 | 0.0011 | 0.692 | 0 | 10 |
| *BRCA1** | 17 | 41201196 | A | G | nonsynonymous SNV | 0.0018 | 0 | 0.625 | 0 | 5 |

| | | | | | | | | | | |
|---|---|---|---|---|---|---|---|---|---|---|
| | 17 | 41223091 | G | A | nonsynonymous SNV | 0.0023 | 0 | 0.544 | 0 | 12 |
| | 17 | 41223240 | A | G | nonsynonymous SNV | 0.0023 | 0 | 0.554 | 0 | 3 |
| | 17 | 41223249 | G | A | nonsynonymous SNV | 0.0038 | 0 | 0.645 | 0 | 22 |
| | 17 | 41226423 | C | T | nonsynonymous SNV | 0.0047 | 4.48E-05 | 0.585 | 0 | 21 |
| | 17 | 41226488 | C | A | nonsynonymous SNV | 6.00E-04 | 0.0037 | 0.516 | 0 | 79 |
| | 17 | 41243509 | T | C | nonsynonymous SNV | 0.001 | 0.0069 | 0.527 | 3 | 94 |
| | 17 | 41246121 | T | C | nonsynonymous SNV | 0.0025 | 0 | 0.576 | 0 | 8 |
| | 17 | 41249297 | G | T | nonsynonymous SNV | 0.0094 | 3.58E-05 | 0.776 | 0 | 46 |
| *BRCA2** | 13 | 32930633 | C | T | nonsynonymous SNV | 0.0034 | 4.48E-05 | 0.519 | 0 | 12 |
| | 13 | 32937488 | G | T | nonsynonymous SNV | 4.00E-04 | 0.0018 | 0.535 | 0 | 30 |
| | 13 | 32972626 | A | T | stopgain | 0.0012 | 0.0089 | NA | 0 | 125 |
| *BTN2A1* | 6 | 26458867 | G | A | nonsynonymous SNV | 6.53E-05 | 0.0023 | 0.527 | 1 | 55 |
| *CCDC60* | 12 | 119978425 | C | T | stopgain | 0.0075 | 5.39E-05 | NA | 0 | 35 |
| *CCDC74A* | 2 | 132285720 | C | G | nonsynonymous SNV | 0.0033 | 9.14E-06 | 0.566 | 0 | 11 |
| | 2 | 132285894 | - | CTGG | frameshift insertion | 0.0828 | 1.00E-04 | NA | 4 | 351 |
| *CENPF* | 1 | 214795521 | T | C | nonsynonymous SNV | 6.54E-05 | 0.0012 | 0.621 | 0 | 13 |
| *CES5A** | 16 | 55899972 | A | T | nonsynonymous SNV | 3.00E-04 | 0.0018 | 0.698 | 0 | 46 |
| | 16 | 55903548 | G | A | nonsynonymous SNV | 0.0491 | 3.00E-04 | 0.538 | 2 | 255 |
| | 16 | 55903554 | G | A | stopgain | 0.1025 | 4.00E-04 | NA | 20 | 465 |
| | 16 | 55905676 | C | T | splicing | 0.0268 | 0.1114 | NA | 120 | 2009 |
| | 16 | 55905677 | T | - | splicing | 3.00E-04 | 0.0011 | NA | 0 | 26 |
| *CFTR** | 7 | 117144344 | C | T | nonsynonymous SNV | 3.00E-04 | 0.0015 | 0.669 | 0 | 47 |
| | 7 | 117149147 | G | A | nonsynonymous SNV | 0.0059 | 0.0285 | 0.677 | 6 | 569 |
| | 7 | 117171029 | G | A | nonsynonymous SNV | 4.00E-04 | 0.0026 | 0.807 | 0 | 37 |
| | 7 | 117171122 | T | C | nonsynonymous SNV | 2.00E-04 | 0.0015 | 0.566 | 0 | 39 |
| | 7 | 117175372 | A | G | nonsynonymous SNV | 0 | 0.0011 | 0.546 | 0 | 7 |
| | 7 | 117176711 | A | T | nonsynonymous SNV | 0.0045 | 0 | 0.917 | 0 | 20 |
| | 7 | 117180174 | G | A | nonsynonymous SNV | 2.00E-04 | 0.0011 | 0.717 | 0 | 13 |
| | 7 | 117199648 | T | G | nonsynonymous SNV | 4.00E-04 | 0.0018 | 0.865 | 0 | 19 |
| | 7 | 117230454 | G | C | nonsynonymous SNV | 0.0021 | 0.0078 | 0.665 | 0 | 100 |
| | 7 | 117232223 | C | T | nonsynonymous SNV | 0.0021 | 0.0095 | 0.706 | 0 | 132 |
| | 7 | 117232481 | G | A | nonsynonymous SNV | 2.00E-04 | 0.0024 | 0.518 | 0 | 39 |

| | | | | | | | | | |
|---|---|---|---|---|---|---|---|---|---|
| | 7 | 117243828 | T | C | nonsynonymous SNV | 3.00E-04 | 0.0011 | 0.659 | 0 | 17 |
| | 7 | 117246808 | G | A | splicing | 0.0014 | 8.98E-06 | NA | 0 | 5 |
| | 7 | 117250575 | G | C | nonsynonymous SNV | 3.00E-04 | 0.0025 | 0.625 | 1 | 89 |
| | 7 | 117267592 | G | T | nonsynonymous SNV | 6.54E-05 | 0.0013 | 0.944 | 0 | 34 |
| | 7 | 117267812 | T | G | nonsynonymous SNV | 0.0021 | 0.0073 | 0.531 | 0 | 151 |
| | 7 | 117282582 | G | A | nonsynonymous SNV | 0.0142 | 1.00E-04 | 0.817 | 0 | 66 |
| | 7 | 117307052 | G | A | nonsynonymous SNV | 0.002 | 3.00E-04 | 0.537 | 0 | 13 |
| *CILP* | 15 | 65489347 | C | T | nonsynonymous SNV | 0.0014 | 1.79E-05 | 0.68 | 0 | 12 |
| *CPT1C* | 19 | 50207994 | T | C | nonsynonymous SNV | 0.0084 | 0 | 0.94 | 0 | 35 |
| *CRIPAK* | 4 | 1388530 | - | CA | frameshift insertion | 0.0018 | 0 | NA | 0 | 3 |
| *CTC1* | 17 | 8131548 | G | A | stopgain | 0.0039 | 5.37E-05 | NA | 0 | 24 |
| | 17 | 8141897 | C | G | nonsynonymous SNV | 9.00E-04 | 0.01 | 0.533 | 0 | 131 |
| *CYP2D6* | 22 | 42522721 | C | T | nonsynonymous SNV | 0.0022 | 1.46E-05 | 0.675 | 0 | 6 |
| | 22 | 42523532 | - | CA | frameshift insertion | 0.0026 | 8.98E-06 | NA | 1 | 15 |
| | 22 | 42523592 | G | A | stopgain | 0.0012 | 5.38E-05 | NA | 0 | 5 |
| | 22 | 42524947 | C | T | splicing | 0.084 | 0.1969 | NA | 406 | 3055 |
| | 22 | 42526694 | G | A | nonsynonymous SNV | 0.1223 | 0.2169 | 0.727 | 612 | 3381 |
| *DDX60L* | 4 | 169305816 | G | A | stopgain | 0.0045 | 4.00E-04 | NA | 1 | 38 |
| | 4 | 169343061 | C | T | splicing | 0.0173 | 0 | NA | 0 | 65 |
| | 4 | 169382849 | C | A | splicing | 0.0018 | 1.35E-05 | NA | 0 | 11 |
| *DNAH6* | 2 | 84881000 | T | C | splicing | 0.003 | 0 | NA | 0 | 7 |
| | 2 | 84932653 | T | G | nonsynonymous SNV | 0.0011 | 0 | 0.82 | 0 | 5 |
| | 2 | 84932866 | G | A | nonsynonymous SNV | 0.0812 | 2.00E-04 | 0.5 | 23 | 307 |
| | 2 | 84934671 | G | A | nonsynonymous SNV | 0.0054 | 0.0012 | 0.7 | 0 | 33 |
| *DNHD1* | 11 | 6555244 | G | T | stopgain | 4.00E-04 | 0.0034 | NA | 1 | 63 |
| | 11 | 6566679 | C | T | stopgain | 0.0014 | 0 | NA | 0 | 7 |
| | 11 | 6567906 | - | AT | frameshift insertion | 0.0105 | 1.00E-04 | NA | 0 | 62 |
| | 11 | 6567907 | - | CCCTAC TGCA | frameshift insertion | 0.3835 | 0.0189 | NA | 0 | 56 |
| | 11 | 6579307 | C | T | stopgain | 3.00E-04 | 0.0016 | NA | 0 | 28 |
| | 11 | 6580337 | G | A | splicing | 0.0016 | 0 | NA | 0 | 7 |
| | 11 | 6585031 | C | T | stopgain | 0.008 | 0 | NA | 0 | 36 |
| | 11 | 6587806 | GTTAC CCCAA | - | splicing | 0.0012 | 0 | NA | 0 | 4 |
| | 11 | 6589683 | G | A | splicing | 0.0033 | 4.48E-05 | NA | 0 | 18 |
| | 11 | 6592054 | C | T | stopgain | 0.005 | 0 | NA | 0 | 12 |

| | | | | | | | | | | |
|---|---|---|---|---|---|---|---|---|---|---|
| *DOCK6* | 19 | 11326148 | - | T | splicing | 0.1467 | 7.00E-04 | NA | 45 | 600 |
| *EFCAB5* | 17 | 28268857 | G | A | splicing | 7.25E-05 | 0.0067 | NA | 1 | 135 |
| | 17 | 28320335 | C | A | stopgain | 2.00E-04 | 0.0014 | NA | 0 | 12 |
| | 17 | 28405349 | C | T | stopgain | 0.0109 | 1.00E-04 | NA | 0 | 50 |
| | 17 | 28407223 | T | C | nonsynonymous SNV | 6.00E-04 | 0.0043 | 0.538 | 0 | 39 |
| *EFCAB6* | 22 | 43950759 | G | A | nonsynonymous SNV | 0.0022 | 6.27E-05 | 0.561 | 0 | 8 |
| *EPPK1** | 8 | 144940779 | - | A | frameshift insertion | 0.1242 | 0.102 | NA | 0 | 2633 |
| | 8 | 144940781 | - | CCATCTC | frameshift insertion | 0.0169 | 0.0432 | NA | 0 | 54 |
| | 8 | 144940792 | - | C | frameshift insertion | 0.0268 | 0.04 | NA | 0 | 10 |
| | 8 | 144940795 | - | G | frameshift insertion | 0.0046 | 0.0055 | NA | 0 | 27 |
| | 8 | 144942300 | C | T | nonsynonymous SNV | 0.0018 | 0.0059 | 0.759 | 0 | 120 |
| | 8 | 144942387 | G | A | nonsynonymous SNV | 3.00E-04 | 0.0034 | 0.805 | 0 | 63 |
| | 8 | 144945092 | G | T | nonsynonymous SNV | 0.0014 | 0 | 0.557 | 0 | 5 |
| | 8 | 144945191 | T | C | nonsynonymous SNV | 0.0388 | 0.1683 | 0.611 | 215 | 2552 |
| | 8 | 144945276 | C | T | nonsynonymous SNV | 0.002 | 0 | 0.584 | 0 | 9 |
| | 8 | 144946188 | G | A | nonsynonymous SNV | 0.0018 | 0.0132 | 0.595 | 2 | 187 |
| | 8 | 144947117 | C | A | nonsynonymous SNV | 4.00E-04 | 0.0037 | 0.626 | 0 | 48 |
| | 8 | 144947129 | G | A | nonsynonymous SNV | 0.002 | 9.36E-06 | 0.726 | 0 | 9 |
| *FER1L6** | 8 | 124978406 | T | A | nonsynonymous SNV | 0.0013 | 0.0087 | 0.831 | 1 | 146 |
| | 8 | 125033892 | C | T | nonsynonymous SNV | 0.004 | 0 | 0.56 | 0 | 18 |
| | 8 | 125052176 | G | A | nonsynonymous SNV | 0.001 | 0.0055 | 0.573 | 1 | 178 |
| | 8 | 125061950 | G | T | nonsynonymous SNV | 4.00E-04 | 0.0021 | 0.841 | 0 | 34 |
| | 8 | 125072527 | T | G | nonsynonymous SNV | 0.0016 | 0.0081 | 0.85 | 0 | 130 |
| | 8 | 125115420 | G | A | nonsynonymous SNV | 0.0126 | 0.0732 | 0.7 | 56 | 1331 |
| *FLG2** | 1 | 152323087 | C | G | stoploss | 0.0057 | 9.13E-06 | NA | 0 | 31 |
| | 1 | 152323132 | G | T | stopgain | 0.27 | 0.1569 | NA | 510 | 3553 |
| | 1 | 152326321 | - | TA | frameshift insertion | 4.00E-04 | 0.0024 | NA | 0 | 36 |
| *FNDC4* | 2 | 27717516 | - | G | frameshift insertion | 0.0025 | 9.00E-04 | NA | 0 | 31 |
| *GALC** | 14 | 88407888 | A | G | nonsynonymous SNV | 0.6115 | 0.4757 | 0.506 | 2836 | 5706 |
| | 14 | 88414158 | G | C | nonsynonymous SNV | 0.0159 | 2.70E-05 | 0.722 | 2 | 67 |
| | 14 | 88431885 | C | T | nonsynonymous SNV | 0.0073 | 8.99E-06 | 0.684 | 0 | 35 |
| | 14 | 88442712 | C | T | nonsynonymous SNV | 0.0371 | 0.1546 | 0.623 | 183 | 2412 |

| | | | | | | | | | | |
|---|---|---|---|---|---|---|---|---|---|---|
| | 14 | 88452926 | T | C | nonsynonymous SNV | 0.0012 | 1.80E-05 | 0.925 | 0 | 5 |
| | 14 | 88452941 | T | C | nonsynonymous SNV | 5.00E-04 | 0.0041 | 0.898 | 0 | 71 |
| | 14 | 88459448 | C | G | nonsynonymous SNV | 0.0327 | 0.1384 | 0.505 | 172 | 2378 |
| *GHDC* | 17 | 40342211 | G | A | stopgain | 0.0015 | 8.98E-06 | NA | 0 | 3 |
| *HEATR4* | 14 | 73965778 | G | A | stopgain | 0.0346 | 3.58E-05 | NA | 1 | 155 |
| *IQCE* | 7 | 2646975 | - | C | frameshift insertion | 3.00E-04 | 0.0036 | NA | 0 | 49 |
| *KCNH2* | 7 | 150644428 | C | A | nonsynonymous SNV | 0.0039 | 0.0228 | 0.6 | 8 | 335 |
| | 7 | 150647283 | G | A | nonsynonymous SNV | 0.0012 | 0 | 0.881 | 0 | 7 |
| | 7 | 150654468 | G | A | nonsynonymous SNV | 2.00E-04 | 0.0013 | 0.545 | 0 | 18 |
| *KIAA1755* | 20 | 36841839 | G | A | stopgain | 0.0128 | 5.06E-05 | NA | 0 | 53 |
| | 20 | 36869005 | G | A | stopgain | 0.0023 | 0.0191 | NA | 0 | 356 |
| *LECT1** | 13 | 53298212 | T | G | nonsynonymous SNV | 5.00E-04 | 0.0034 | 0.617 | 0 | 26 |
| *LIPE* | 19 | 42914668 | G | A | nonsynonymous SNV | 0.0011 | 0.0053 | 0.566 | 0 | 103 |
| | 19 | 42930751 | G | T | stopgain | 0.0032 | 3.58E-05 | NA | 0 | 13 |
| *MICALL2* | 7 | 1488319 | C | G | nonsynonymous SNV | 0.0011 | 0 | 0.796 | 0 | 3 |
| *MYBPC3* | 11 | 47353755 | G | A | nonsynonymous SNV | 0.0014 | 2.00E-04 | 0.607 | 0 | 14 |
| | 11 | 47355294 | G | A | nonsynonymous SNV | 0.0076 | 3.75E-05 | 0.628 | 0 | 38 |
| | 11 | 47364234 | C | T | nonsynonymous SNV | 0.0053 | 2.00E-04 | 0.858 | 0 | 25 |
| *MYCBP2* | 13 | 77831927 | C | G | nonsynonymous SNV | 0.0021 | 0 | 0.546 | 0 | 5 |
| *MYO1A* | 12 | 57424918 | G | A | nonsynonymous SNV | 7.00E-04 | 0.007 | 0.674 | 0 | 153 |
| | 12 | 57430138 | G | A | nonsynonymous SNV | 1.00E-04 | 0.0012 | 0.581 | 0 | 14 |
| | 12 | 57431355 | T | A | nonsynonymous SNV | 7.00E-04 | 0.0026 | 0.837 | 0 | 62 |
| | 12 | 57431732 | G | C | nonsynonymous SNV | 6.54E-05 | 0.0011 | 0.82 | 1 | 59 |
| | 12 | 57437952 | G | C | nonsynonymous SNV | 0.0043 | 8.96E-06 | 0.66 | 0 | 16 |
| | 12 | 57441459 | G | A | stopgain | 6.00E-04 | 0.0042 | NA | 0 | 75 |
| | 12 | 57441501 | C | A | nonsynonymous SNV | 3.00E-04 | 0.0024 | 0.736 | 0 | 31 |
| | 12 | 57442059 | C | G | nonsynonymous SNV | 0.0106 | 4.50E-05 | 0.552 | 0 | 52 |
| *PIWIL3* | 22 | 25124022 | TTAAGT | - | splicing | 0.0075 | 0 | NA | 0 | 39 |
| | 22 | 25124143 | C | T | splicing | 0.0024 | 0.0081 | NA | 2 | 120 |
| | 22 | 25130050 | C | A | splicing | 0.0016 | 0 | NA | 0 | 5 |
| *PLSCR5* | 3 | 146307508 | C | T | nonsynonymous SNV | 0.0011 | 3.61E-05 | 0.517 | 0 | 6 |
| *POLN** | 4 | 2087404 | A | C | nonsynonymous SNV | 0.0234 | 3.58E-05 | 0.802 | 0 | 96 |
| | 4 | 2097595 | G | A | nonsynonymous SNV | 0.0019 | 4.48E-05 | 0.625 | 0 | 7 |

| | | | | | | | | | |
|---|---|---|---|---|---|---|---|---|---|
| | 4 | 2129933 | C | T | nonsynonymous SNV | 0.0016 | 0 | 0.803 | 0 | 12 |
| | 4 | 2160884 | G | A | stopgain | 0.0025 | 0 | NA | 0 | 8 |
| *PSG8* | 19 | 43258740 | C | G | splicing | 0.0043 | 9.17E-06 | NA | 0 | 19 |
| *RNPEP* | 1 | 201965367 | T | C | nonsynonymous SNV | 0.0012 | 0 | 0.542 | 0 | 7 |
| *RTKN2** | 10 | 63959584 | A | G | nonsynonymous SNV | 0.0016 | 1.79E-05 | 0.817 | 0 | 6 |
| | 10 | 63976968 | C | T | nonsynonymous SNV | 6.00E-04 | 0.0019 | 0.6 | 0 | 36 |
| | 10 | 64022582 | - | A | splicing | 0.0039 | 0.0062 | NA | 0 | 68 |
| *SCNN1D** | 1 | 1219382 | C | T | stopgain | 0.0103 | 7.79E-05 | 0.203 | 0 | 35 |
| | 1 | 1222519 | C | T | stopgain | 0.0034 | 2.75E-05 | NA | 0 | 13 |
| *SEC14L3* | 22 | 30860831 | G | A | nonsynonymous SNV | 0.0097 | 1.79E-05 | 0.641 | 0 | 37 |
| | 22 | 30864621 | C | T | stopgain | 0.0014 | 0 | 0.709 | 0 | 1 |
| | 22 | 30864655 | A | G | nonsynonymous SNV | 0.0022 | 0.0206 | 0.799 | 5 | 336 |
| | 22 | 30866053 | G | A | nonsynonymous SNV | 0.0022 | 0.0204 | 0.797 | 5 | 336 |
| | 22 | 30866240 | G | A | nonsynonymous SNV | 0.0015 | 2.71E-05 | 0.747 | 0 | 3 |
| | 22 | 30866506 | G | A | nonsynonymous SNV | 0.0017 | 8.95E-06 | 0.924 | 0 | 10 |
| *SLC12A3* | 16 | 56899184 | G | C | nonsynonymous SNV | 3.00E-04 | 0.0011 | 0.509 | 0 | 16 |
| *SLC22A2* | 6 | 160668263 | T | A | stopgain | 0.0012 | 0 | NA | 0 | 5 |
| *SLC2A8* | 9 | 130166296 | C | T | nonsynonymous SNV | 0.0023 | 5.42E-05 | 0.721 | 0 | 9 |
| | 9 | 130167787 | C | G | nonsynonymous SNV | 0.0125 | 1.80E-05 | 0.512 | 0 | 51 |
| | 9 | 130167800 | T | C | nonsynonymous SNV | 0.0034 | 8.99E-06 | 0.854 | 0 | 12 |
| *SPG7* | 16 | 89598369 | G | A | nonsynonymous SNV | 2.00E-04 | 0.0016 | 0.948 | 0 | 11 |
| | 16 | 89613145 | C | T | nonsynonymous SNV | 0.0012 | 0.0049 | 0.923 | 1 | 122 |
| *STOX2* | 4 | 184930308 | CAG | - | splicing | 9.00E-04 | 0.0026 | NA | 0 | 1 |
| *SYT15* | 10 | 46962104 | G | A | stopgain | 0.0021 | 2.73E-05 | NA | 0 | 8 |
| | 10 | 46962114 | T | C | splicing | 4.00E-04 | 0.0014 | NA | 0 | 27 |
| | 10 | 46964021 | T | C | splicing | 0.0026 | 0 | NA | 0 | 12 |
| | 10 | 46965803 | C | T | nonsynonymous SNV | 3.00E-04 | 0.0021 | 0.875 | 0 | 72 |
| | 10 | 46965887 | - | G | splicing | 0.073 | 0.2696 | NA | 1 | 5462 |
| *TGM6** | 20 | 2376062 | T | A | nonsynonymous SNV | 0.0012 | 0 | 0.893 | 0 | 6 |
| | 20 | 2380277 | C | T | nonsynonymous SNV | 0.0012 | 4.50E-05 | 0.955 | 0 | 4 |
| | 20 | 2380285 | T | C | nonsynonymous SNV | 0.0029 | 0 | 0.925 | 0 | 10 |
| | 20 | 2384078 | G | A | nonsynonymous SNV | 0.0018 | 8.95E-06 | 0.874 | 0 | 7 |
| | 20 | 2384241 | G | A | nonsynonymous SNV | 0.007 | 8.96E-06 | 0.925 | 1 | 15 |
| | 20 | 2384259 | G | A | nonsynonymous SNV | 0.024 | 1.00E-04 | 0.813 | 4 | 105 |

| | | | | | | | | | | |
|---|---|---|---|---|---|---|---|---|---|---|
| | 20 | 2398064 | G | A | nonsynonymous SNV | 6.55E-05 | 0.0023 | 0.853 | 0 | 44 |
| | 20 | 2398097 | A | C | nonsynonymous SNV | 0.0011 | 0 | 0.68 | 0 | 3 |
| | 20 | 2411118 | T | A | nonsynonymous SNV | 0.002 | 0 | 0.637 | 0 | 7 |
| *TRDN* | 6 | 123594510 | - | A | splicing | 0.0864 | 0.1639 | NA | 369 | 3116 |
| | 6 | 123673672 | - | T | stoploss | 0.0065 | 4.00E-04 | NA | 0 | 21 |
| | 6 | 123786032 | - | A | frameshift insertion | 0.0017 | 0.0035 | NA | 0 | 348 |
| *TTC21B* | 2 | 166773990 | A | C | nonsynonymous SNV | 0.0056 | 0 | 0.716 | 0 | 33 |
| | 2 | 166775889 | G | C | nonsynonymous SNV | 0.012 | 0 | 0.551 | 0 | 45 |
| *TTN** | 2 | 179391962 | C | T | nonsynonymous SNV | 0.0011 | 0 | 0.594 | 0 | 8 |
| | 2 | 179396215 | G | A | nonsynonymous SNV | 0.0061 | 3.59E-05 | 0.523 | 0 | 23 |
| | 2 | 179414177 | G | A | nonsynonymous SNV | 6.57E-05 | 0.0032 | 0.52 | 0 | 61 |
| | 2 | 179422944 | A | C | nonsynonymous SNV | 0.0029 | 0 | 0.551 | 0 | 9 |
| | 2 | 179427560 | G | T | nonsynonymous SNV | 0.0024 | 9.08E-06 | 0.639 | 0 | 12 |
| | 2 | 179429557 | C | A | nonsynonymous SNV | 0.0011 | 0 | 0.631 | 0 | 4 |
| | 2 | 179431076 | C | G | nonsynonymous SNV | 0.0036 | 0.0198 | 0.689 | 1 | 351 |
| | 2 | 179431963 | A | T | nonsynonymous SNV | 0.002 | 0 | 0.608 | 0 | 3 |
| | 2 | 179433046 | C | G | nonsynonymous SNV | 0.0027 | 3.00E-04 | 0.53 | 0 | 26 |
| | 2 | 179438713 | A | G | nonsynonymous SNV | 2.00E-04 | 0.0011 | 0.863 | 0 | 13 |
| | 2 | 179438866 | C | T | nonsynonymous SNV | 0.0787 | 0.0326 | 0.517 | 25 | 1009 |
| | 2 | 179439907 | A | C | nonsynonymous SNV | 0.0085 | 5.45E-05 | 0.51 | 1 | 41 |
| | 2 | 179440163 | C | G | nonsynonymous SNV | 0.002 | 0.0124 | 0.639 | 2 | 227 |
| | 2 | 179441038 | C | T | nonsynonymous SNV | 2.00E-04 | 0.0021 | 0.601 | 0 | 37 |
| | 2 | 179441295 | T | C | nonsynonymous SNV | 0.0012 | 0.0085 | 0.566 | 1 | 219 |
| | 2 | 179441932 | G | A | nonsynonymous SNV | 0.0016 | 0.0055 | 0.593 | 0 | 107 |
| | 2 | 179442784 | C | G | nonsynonymous SNV | 3.00E-04 | 0.0026 | 0.761 | 0 | 61 |
| | 2 | 179447754 | C | T | nonsynonymous SNV | 0.0028 | 3.64E-05 | 0.549 | 0 | 14 |
| | 2 | 179449579 | C | T | nonsynonymous SNV | 2.00E-04 | 0.0016 | 0.518 | 0 | 39 |
| | 2 | 179449606 | C | T | nonsynonymous SNV | 0.0081 | 1.00E-04 | 0.525 | 0 | 38 |
| | 2 | 179466177 | A | G | nonsynonymous SNV | 0.0071 | 0 | 0.505 | 0 | 28 |
| | 2 | 179466400 | T | C | nonsynonymous SNV | 0.0022 | 9.36E-06 | 0.5 | 0 | 6 |
| | 2 | 179472223 | A | G | nonsynonymous SNV | 0.0024 | 0.0139 | 0.801 | 2 | 245 |

| | | | | | | | | | | |
|---|---|---|---|---|---|---|---|---|---|---|
| | 2 | 179479288 | A | G | nonsynonymous SNV | 3.00E-04 | 0.0012 | 0.748 | 0 | 40 |
| | 2 | 179482994 | G | A | nonsynonymous SNV | 7.00E-04 | 0.0037 | 0.56 | 0 | 87 |
| | 2 | 179486345 | T | A | nonsynonymous SNV | 0.0564 | 7.00E-04 | 0.503 | 5 | 265 |
| | 2 | 179563643 | - | A | splicing | 0.0723 | 0.0924 | NA | 34 | 424 |
| | 2 | 179563643 | - | AA | splicing | 0.0021 | 0.0038 | NA | 0 | 3 |
| | 2 | 179571683 | T | G | splicing | 0.0018 | 1.00E-05 | NA | 0 | 9 |
| | 2 | 179585187 | C | T | nonsynonymous SNV | 5.00E-04 | 0.0014 | 0.593 | 0 | 21 |
| | 2 | 179587130 | C | G | nonsynonymous SNV | 0.1722 | 0.15 | 0.532 | 318 | 3030 |
| | 2 | 179588813 | C | T | nonsynonymous SNV | 0.0098 | 5.00E-04 | 0.632 | 0 | 68 |
| | 2 | 179589083 | T | A | nonsynonymous SNV | 0.0165 | 1.80E-05 | 0.586 | 2 | 77 |
| | 2 | 179590329 | C | T | nonsynonymous SNV | 0.0031 | 0.0179 | 0.596 | 4 | 295 |
| | 2 | 179590714 | T | A | nonsynonymous SNV | 3.00E-04 | 0.0012 | 0.501 | 0 | 18 |
| | 2 | 179592567 | G | A | nonsynonymous SNV | 0.0063 | 9.37E-06 | 0.579 | 0 | 26 |
| | 2 | 179598553 | T | G | nonsynonymous SNV | 5.00E-04 | 0.0021 | 0.561 | 0 | 22 |
| | 2 | 179615060 | C | T | nonsynonymous SNV | 0.0104 | 4.00E-04 | 0.76 | 0 | 61 |
| | 2 | 179620947 | A | G | splicing | 0.0042 | 0 | NA | 0 | 24 |
| | 2 | 179628969 | G | A | nonsynonymous SNV | 0.0015 | 7.17E-05 | 0.521 | 0 | 3 |
| | 2 | 179638721 | C | T | nonsynonymous SNV | 0.0936 | 0.0234 | 0.571 | 39 | 898 |
| | 2 | 179639711 | C | A | nonsynonymous SNV | 0.0297 | 3.59E-05 | 0.644 | 0 | 116 |
| | 2 | 179640923 | G | A | nonsynonymous SNV | 0.0017 | 2.00E-04 | 0.626 | 0 | 20 |
| | 2 | 179643775 | C | T | nonsynonymous SNV | 0.0012 | 0.0084 | 0.679 | 1 | 135 |
| | 2 | 179645962 | C | G | nonsynonymous SNV | 3.00E-04 | 0.0035 | 0.71 | 0 | 72 |
| *WDR27** | 6 | 170036466 | C | A | splicing | 0.0877 | 0.0061 | NA | 18 | 460 |
| | 6 | 170060821 | G | C | stopgain | 0.0012 | 0 | NA | 0 | 3 |
| | 6 | 170060843 | G | A | stopgain | 2.00E-04 | 0.0013 | NA | 0 | 26 |
| *WDR87** | 19 | 38379447 | G | A | stopgain | 0 | 0.0015 | NA | 0 | 26 |
| | 19 | 38383925 | - | AATATCTG | frameshift insertion | 0.0017 | 8.64E-05 | NA | 0 | 15 |
| *ZNF266* | 19 | 9524487 | G | A | stopgain | 0.0021 | 2.69E-05 | NA | 0 | 8 |
| *ZNF334* | 20 | 45131134 | G | A | stopgain | 0.0013 | 5.39E-05 | NA | 0 | 8 |

**Table S4**. **Number of carriers for low-frequency to common pLOF and predicted deleterious missense variants among 106 genes of interest.**

List of 106 genes with associations p<E-06 from exome-by-phenome-wide association studies using pLOF-based gene burdens. Each gene is labeled with low frequency to common (minor allele frequency > 0.1% per gnomAD among Europeans and Africans) pLOF variants and missense variants with REVEL scores of at least 0.5 that were prevalent in PMBB. Each variant is labeled with chromosomal position based on build GRCh37/hg19, the type of exonic variant, minor allele frequencies (MAF) specific to Africans (AFR) and Non-Finnish Europeans (NFE) according to gnomAD, REVEL scores if missense, and the number of homozygous and heterozygous carriers in PMBB. Note: starred genes (*) represent those for which univariate studies successfully replicated associations identified from the pLOF-based gene burden studies.

### Table S5

| pLOF-Based Gene Burden | | | | | Single Variants | | | | | | | | | |
|---|---|---|---|---|---|---|---|---|---|---|---|---|---|---|
| Gene | Phecode | Description | OR | P | Chr | Position | Ref | Alt | Phecode | Description | OR | P | N | rs |
| *BCA10* | 514.1 | Abnormal results of function study of pulmonary system | 36.73 | 1.54E-07 | 17 | 67150465 | - | ACCT GGAA | 514.2 | Solitary pulmonary nodule | 1.63 | 2.78E-03 | 217 hom, 1018 het | rs1379626 |
| *AIM1* | 741 | Symptoms and disorders of the joints | 11.29 | 3.05E-07 | 6 | 106991361 | T | C | 741.2 | Stiffness of joint | 2.89 | 6.41E-03 | 37 hom, 1100 het | rs617411 |
| *ASPH* | 681.3 | Cellulitis and abscess of arm/hand | 74.06 | 5.39E-10 | 8 | 62550924 | - | AA | 681.1 | Cellulitis and abscess of fingers/toes | 3.62 | 1.81E-02 | 481 het | NA |
| *BRCA1* | 433.6 | Acute, but ill-defined cerebrovascular disease | 41.19 | 3.21E-08 | 17 | 41226488 | C | A | 433.21 | Cerebral artery occlusion, with cerebral infarction | 2.48 | 4.06E-02 | 79 het | rs18007 |
| | | | | | | | | | 433.2 | Occlusion of cerebral arteries | 2.45 | 4.32E-02 | | |
| *BRCA2* | 368.2 | Diplopia and disorders of binocular vision | 16.48 | 7.99E-07 | 13 | 32937488 | G | T | 368.4 | Visual field defects | 14.28 | 1.16E-02 | 30 het | rs288977 |
| *CES5A* | 286.9 | Abnormal coagulation profile | 27.27 | 8.10E-08 | 16 | 55905677 | T | - | 286.9 | Abnormal coagulation profile | 11.36 | 2.14E-03 | 26 het | NA |
| *CFTR* | 496.3 | Bronchiectasis | 20.99 | 4.21E-11 | 7 | 117149147 | G | A | 496.21 | Obstructive chronic bronchitis | 2.38 | 6.85E-03 | 6 hom, 569 het | rs18000 |
| | | | | | | | | | 496.2 | Chronic bronchitis | 1.74 | 4.33E-02 | | |
| | | | | | | | | | 496.3 | Bronchiectasis | 2.00 | 4.62E-02 | | |
| *EPPK1* | 451.2 | Phlebitis and thrombophlebitis of lower extremities | 14.85 | 9.19E-08 | 8 | 144942300 | C | T | 451.2 | Phlebitis and thrombophlebitis of lower extremities | 5.42 | 6.30E-03 | 120 het | rs1915331 |
| | | | | | | | | | 451 | Phlebitis and thrombophlebitis | 3.34 | 4.74E-02 | | |
| *ER1L6* | 772.1 | Muscular wasting and disuse atrophy | 24.47 | 7.18E-07 | 8 | 125061950 | G | T | 772.1 | Muscular wasting and disuse atrophy | 9.27 | 3.36E-02 | 34 het | rs2008943 |
| *FLG2* | 741.2 | Stiffness of joint | 23.88 | 1.76E-07 | 1 | 152323132 | G | T | 741.2 | Stiffness of joint | 2.70 | 9.65E-04 | 510 hom, 3553 het | rs125687 |
| *GALC* | 596 | Other disorders of bladder | 12.93 | 3.48E-07 | 14 | 88442712* | C | T | 596 | Other disorders of bladder | 1.35 | 3.52E-02 | 183 hom, 2412 het | rs343627 |
| | | | | | 14 | 88459448* | C | G | 596 | Other disorders of bladder | 1.35 | 3.57E-02 | 172 hom, 2378 het | rs111887 |
| *LECT1* | 555.2 | Ulcerative colitis | 27.03 | 7.11E-07 | 13 | 53298212 | T | G | 555 | Inflammatory bowel disease and other gastroenteritis and colitis | 9.14 | 3.45E-03 | 26 het | rs1407346 |

| | | | | | | | | | | | | | | |
|---|---|---|---|---|---|---|---|---|---|---|---|---|---|---|
| | | | | | | | | | 555.1 | Regional enteritis | 10.19 | 2.67E-02 | | |
| *POLN* | 289 | Other diseases of blood and blood-forming organs | 47.40 | 2.57E-08 | 4 | 2129933 | C | T | 289.4 | Lymphadenitis | 10.98 | 2.76E-03 | 12 het | rs1480621 |
| *RTKN2* | 458.1 | Orthostatic hypotension | 84.42 | 7.24E-07 | 10 | 64022582 | - | A | 458.1 | Orthostatic hypotension | 45.55 | 2.19E-04 | 68 het | NA |
| *CNN1D* | 426.8 | Other cardiac conduction disorders | 65.97 | 4.52E-07 | 1 | 1219382 | C | T | 426.24 | Atrioventricular block, complete | 7.42 | 1.66E-02 | 35 het | NA |
| | | | | | | | | | 426.91 | Cardiac pacemaker in situ | 3.79 | 4.43E-02 | | |
| *TGM6* | 214 | Lipoma | 13.08 | 2.77E-07 | 20 | 2384259 | G | A | 214 | Lipoma | 3.09 | 9.81E-03 | 4 hom, 105 het | rs797240 |
| *TTN* | 425 | Cardiomyopathy | 2.40 | 7.83E-13 | 2 | 179431076 | C | G | 426.21 | First degree AV block | 2.26 | 4.21E-02 | 1 hom, 351 het | rs563072 |
| | 425.1 | Primary/intrinsic cardiomyopathies | 2.40 | 1.11E-11 | | | | | 427.6 | Premature beats | 0.59 | 4.72E-02 | | |
| | 426.92 | Cardiac defibrillator in situ | 2.69 | 1.77E-08 | 2 | 179433046 | C | G | 426.21 | First degree AV block | 29.65 | 2.13E-02 | 26 het | rs1866811 |
| | 427.1 | Paroxysmal tachycardia, unspecified | 2.57 | 6.45E-09 | | | | | 426.3 | Bundle branch block | 4.43 | 4.60E-02 | | |
| | 427.6 | Premature beats | 3.04 | 8.90E-09 | | | | | 427.5 | Arrhythmia (cardiac) NOS | 6.18 | 3.87E-02 | | |
| | 427.12 | Paroxysmal ventricular tachycardia | 2.60 | 2.18E-08 | 2 | 179440163 | C | G | 425.1 | Primary/intrinsic cardiomyopathies | 1.47 | 2.81E-02 | 2 hom, 227 het | rs558011 |
| | | | | | | | | | 426.9 | Cardiac pacemaker/device in situ | 1.68 | 1.14E-02 | | |
| | | | | | | | | | 426.92 | Cardiac defibrillator in situ | 1.73 | 1.43E-02 | | |
| | | | | | | | | | 426.24 | Atrioventricular block, complete | 2.17 | 2.26E-02 | | |
| | | | | | | | | | 426.2 | Atrioventricular [AV] block | 1.93 | 2.69E-02 | | |
| | | | | | | | | | 427.6 | Premature beats | 1.94 | 6.47E-03 | | |
| | | | | | | | | | 427.9 | Palpitations | 1.68 | 8.24E-03 | | |
| | | | | | | | | | 427.5 | Arrhythmia (cardiac) NOS | 2.76 | 2.33E-02 | | |
| | | | | | | | | | 427.8 | Sinoatrial node dysfunction (Bradycardia) | 1.85 | 3.97E-02 | | |
| | | | | | | | | | 427.22 | Atrial flutter | 1.60 | 4.77E-02 | | |

| | | | | | | | | | |
|---|---|---|---|---|---|---|---|---|---|
| 2 | 179441038 | C | T | 426.9 | Cardiac pacemaker/device in situ | 0.10 | 2.85E-02 | 37 het | rs2010439 |
| | | | | 427.2 | Atrial fibrillation and flutter | 0.23 | 5.96E-03 | | |
| | | | | 427.21 | Atrial fibrillation | 0.25 | 7.85E-03 | | |
| | | | | 427 | Cardiac dysrhythmias | 0.34 | 7.90E-03 | | |
| | | | | 427.1 | Paroxysmal tachycardia, unspecified | 0.10 | 2.75E-02 | | |
| 2 | 179441932 | G | A | 426.24 | Atrioventricular block, complete | 3.51 | 2.19E-03 | 107 het | rs559804 |
| | | | | 426.22 | Mobitz II AV block | 5.73 | 2.42E-02 | | |
| | | | | 426.23 | Second degree AV block | 5.71 | 2.48E-02 | | |
| 2 | 179449579 | C | T | 426.25 | Other heart block | 18.26 | 2.09E-03 | 39 het | rs1506619 |
| 2 | 179479288 | A | G | 426.4 | Anomalous atrioventricular excitation | 11.54 | 3.16E-02 | 40 het | rs726772 |
| | | | | 427.7 | Tachycardia NOS | 6.34 | 7.14E-03 | | |
| 2 | 179563643 | - | A | 426 | Cardiac conduction disorders | 0.75 | 3.32E-02 | 34 het | NA |
| | | | | 426.9 | Cardiac pacemaker/device in situ | 0.74 | 4.74E-02 | | |
| 2 | 179585187 | C | T | 426.8 | Other cardiac conduction disorders | 15.87 | 1.74E-02 | 21 het | rs726489 |
| | | | | 427.42 | Cardiac arrest | 7.33 | 2.27E-02 | | |
| 2 | 179587130 | C | G | 425.2 | Secondary/extrinsic cardiomyopathies | 0.80 | 9.10E-03 | 318 hom, 3030 het | rs126931 |
| | | | | 425.1 | Primary/intrinsic cardiomyopathies | 0.88 | 1.74E-02 | | |
| | | | | 426.92 | Cardiac defibrillator in situ | 0.81 | 2.28E-03 | | |
| | | | | 426.24 | Atrioventricular block, complete | 0.70 | 5.15E-03 | | |
| | | | | 426.9 | Cardiac pacemaker/device in situ | 0.86 | 1.06E-02 | | |
| | | | | 426.2 | Atrioventricular [AV] block | 0.81 | 2.82E-02 | | |

| | | | | | | | | | |
|---|---|---|---|---|---|---|---|---|---|
| 2 | 179588813 | C | T | 425.11 | Hypertrophic obstructive cardiomyopathy | 7.54 | 9.71E-04 | 68 het | rs726489 |
| | | | | 425.2 | Secondary/extrinsic cardiomyopathies | 2.87 | 4.84E-02 | | |
| | | | | 426.7 | Abnormal electrocardiogram [ECG] [EKG] | 3.55 | 1.04E-02 | | |
| | | | | 426.22 | Mobitz II AV block | 5.79 | 4.13E-02 | | |
| 2 | 179590329 | C | T | 426.4 | Anomalous atrioventricular excitation | 4.52 | 4.35E-03 | 4 hom, 295 het | rs173554 |
| | | | | 426.3 | Bundle branch block | 1.91 | 5.38E-03 | | |
| | | | | 426.31 | Right bundle branch block | 2.33 | 7.68E-03 | | |
| | | | | 427.41 | Ventricular fibrillation and flutter | 2.58 | 2.03E-02 | | rs173554 |
| | | | | 427.6 | Premature beats | 1.61 | 2.44E-02 | | rs173554 |
| | | | | 427.21 | Atrial fibrillation | 1.41 | 2.81E-02 | | rs173554 |
| | | | | 427.2 | Atrial fibrillation and flutter | 1.40 | 2.94E-02 | | rs173554 |
| | | | | 427.61 | Supraventricular premature beats | 2.20 | 4.76E-02 | | rs173554 |
| 2 | 179598553 | T | G | 426 | Cardiac conduction disorders | 0.27 | 4.77E-02 | 22 het | rs726489 |
| | | | | 427 | Cardiac dysrhythmias | 0.25 | 6.39E-03 | | |
| | | | | 427.21 | Atrial fibrillation | 0.18 | 8.12E-03 | | |
| | | | | 427.2 | Atrial fibrillation and flutter | 0.29 | 2.43E-02 | | |
| | | | | 427.1 | Paroxysmal tachycardia, unspecified | 0.12 | 4.28E-02 | | |
| 2 | 179615060 | C | T | 427.5 | Arrhythmia (cardiac) NOS | 24.86 | 2.16E-04 | 61 het | rs143253[illegible] |
| 2 | 179638721 | C | T | 426.31 | Right bundle branch block | 0.47 | 3.36E-02 | 39 hom, 898 het | rs48940[illegible] |
| 2 | 179640923 | G | A | 426.25 | Other heart block | 31.95 | 6.08E-03 | 20 het | rs146496[illegible] |
| | | | | 426.24 | Atrioventricular block, complete | 6.14 | 3.61E-02 | | |
| 2 | 179645962 | C | G | 425 | Cardiomyopathy | 1.90 | 3.03E-02 | 72 het | rs726478 |

| | | | | | | | | | | | | | | |
|---|---|---|---|---|---|---|---|---|---|---|---|---|---|---|
| | | | | | | | | | 426.2 | Atrioventricular [AV] block | 2.68 | 3.73E-02 | | |
| | | | | | | | | | 427.8 | Sinoatrial node dysfunction (Bradycardia) | 3.31 | 6.98E-03 | | |
| | | | | | | | | | 427.6 | Premature beats | 2.29 | 4.32E-02 | | |
| *VDR27* | 695.7 | Prurigo and Lichen | 64.57 | 6.31E-07 | 6 | 170060843 | G | A | 695.8 | Other specified erythematous conditions | 12.61 | 1.62E-02 | 26 het | rs2011228 |
| *VDR87* | 550.5 | Ventral hernia | 8.34 | 1.70E-07 | 19 | 38379447 | G | A | 550.4 | Umbilical hernia | 26.95 | 3.02E-05 | 26 het | rs1897043 |
| | | | | | | | | | 550.6 | Incisional hernia | 7.55 | 9.66E-03 | | |
| | | | | | | | | | 550.5 | Ventral hernia | 8.40 | 4.33E-02 | | |
| | | | | | 19 | 38383925 | - | AATA TCTG | 550.5 | Ventral hernia | 214.40 | 3.04E-04 | 15 het | NA |

*variants in LD with each other, but not with variants included in pLOF burden

**Table S5: Evaluation of robustness via univariate association studies within Penn Medicine Biobank.**

List of significantly replicated associations via univariate analyses of low-frequency to common (MAF > 0.1%) predicted loss-of-function (pLOF) or predicted deleterious missense (REVEL ≥ 0.5) variants in Penn Medicine Biobank (PMBB) among 106 genes for which predicted loss-of-function (pLOF)-based gene burdens had p<E-06 from exome-by-phenome-wide association studies in PMBB. Summary statistics are based on an exact logistic regression model adjusted for age, $age^2$, gender, and the first ten principal components of ancestry. Reported are the odds ratios and p-values resulting from meta-analysis of the summary statistics obtained from running the logistic regression model separately in individuals of European and African ancestry, and the number of carriers for each gene burden or single variant. Additionally, single variants are annotated by their genomic location according to GRCh37/hg19, as well as their rs identification codes.

### Table S6

| Phecode | Description | PMBB | | | UKBB | | | p |
|---|---|---|---|---|---|---|---|---|
| | | Cases | Controls | Case Prevalence | Cases | Controls | Case Prevalence | |
| 8.6 | Viral Enteritis | 45 | 9786 | 0.458% | 74 | 34305 | 0.215% | 6.84E-05 |
| 31 | Diseases due to other mycobacteria | 22 | 6982 | 0.314% | 5 | 33255 | 0.015% | 1.42E-17 |
| 130 | Spirochetal infection | 47 | 7787 | 0.600% | 6 | 34583 | 0.017% | 1.15E-38 |
| 130.1 | Lyme disease | 46 | 7787 | 0.587% | 3 | 34583 | 0.009% | 4.13E-41 |
| 145 | Cancer of mouth | 44 | 8004 | 0.547% | 64 | 34425 | 0.186% | 1.39E-08 |
| 145.2 | Cancer of tongue | 27 | 8004 | 0.336% | 34 | 34425 | 0.099% | 9.62E-07 |
| 174 | Breast cancer | 247 | 9659 | 2.493% | 1230 | 32678 | 3.627% | 4.50E-08 |
| 174.1 | Breast cancer [female] | 246 | 9659 | 2.484% | 1224 | 32678 | 3.610% | 5.13E-08 |
| 174.11 | Malignant neoplasm of female breast | 235 | 9659 | 2.375% | 1147 | 32678 | 3.391% | 4.49E-07 |
| 198.1 | Secondary malignancy of lymph nodes | 52 | 7417 | 0.696% | 332 | 31072 | 1.057% | 5.60E-03 |
| 198.4 | Secondary malignant neoplasm of liver | 30 | 7417 | 0.403% | 185 | 31072 | 0.592% | 5.94E-02 |
| 201 | Hodgkin's disease | 52 | 7802 | 0.662% | 24 | 34238 | 0.070% | 3.70E-28 |
| 214 | Lipoma | 89 | 9993 | 0.883% | 715 | 33804 | 2.071% | 4.20E-15 |
| 225 | Benign neoplasm of brain and other parts of nervous system | 39 | 8020 | 0.484% | 70 | 34483 | 0.203% | 1.19E-05 |
| 225.1 | Benign neoplasm of brain, cranial nerves, meninges | 36 | 8020 | 0.447% | 68 | 34483 | 0.197% | 7.17E-05 |
| 246.7 | Abnormal results of function study of thyroid | 55 | 6266 | 0.870% | 5 | 32881 | 0.015% | 6.95E-56 |
| 250.12 | Type 1 diabetes with renal manifestations | 49 | 6143 | 0.791% | 1 | 32689 | 0.003% | 2.16E-55 |
| 250.13 | Type 1 diabetes with ophthalmic manifestations | 23 | 5198 | 0.441% | 33 | 32689 | 0.101% | 9.29E-09 |
| 251 | Other disorders of pancreatic internal secretion | 42 | 7502 | 0.557% | 1 | 34254 | 0.003% | 7.37E-41 |
| 260.6 | Anorexia | 60 | 8276 | 0.720% | 69 | 34400 | 0.200% | 1.93E-14 |
| 271 | Disorders of carbohydrate transport and metabolism | 21 | 8082 | 0.259% | 19 | 34610 | 0.055% | 1.87E-07 |
| 274 | Gout and other crystal arthropathies | 528 | 7384 | 6.673% | 388 | 34241 | 1.120% | 2.00E-206 |
| 274.1 | Gout | 521 | 7384 | 6.591% | 318 | 34241 | 0.920% | 1.20E-233 |
| 274.11 | Gouty arthropathy | 186 | 9210 | 1.980% | 5 | 34241 | 0.015% | 4.01E-143 |
| 281 | Other deficiency anemia | 62 | 6148 | 0.998% | 102 | 32782 | 0.310% | 3.23E-14 |
| 286.9 | Abnormal coagulation profile | 156 | 6106 | 2.491% | NA** | NA** | NA** | NA |
| 289 | Other diseases of blood and blood-forming organs | 32 | 6962 | 0.458% | 96 | 33977 | 0.282% | 2.23E-02 |
| 296.1 | Bipolar | 146 | 6939 | 2.061% | 115 | 31247 | 0.367% | 6.89E-55 |
| 300.12 | Agorophobia, social phobia, and panic disorder | 78 | 6939 | 1.112% | 68 | 31247 | 0.217% | 1.33E-27 |
| 334 | Degenerative disease of the spinal cord | 35 | 7286 | 0.478% | 143 | 33472 | 0.425% | 6.01E-01 |
| 345.1 | Epilepsy | 68 | 9071 | 0.744% | 79 | 33472 | 0.235% | 3.94E-13 |

| | | | | | | | | |
|---|---|---|---|---|---|---|---|---|
| 345.12 | Partial epilepsy | 52 | 9071 | 0.570% | 19 | 33472 | 0.057% | 7.66E-26 |
| 350.3 | Lack of coordination | 28 | 7578 | 0.368% | 33 | 34406 | 0.096% | 4.24E-08 |
| 356 | Hereditary and idiopathic peripheral neuropathy | 164 | 9642 | 1.672% | 8 | 34429 | 0.023% | 1.13E-117 |
| 359 | Muscular dystrophies and other myopathies | 25 | 7701 | 0.324% | 53 | 34429 | 0.154% | 2.74E-03 |
| 359.2 | Myopathy | 20 | 7701 | 0.259% | 43 | 34429 | 0.125% | 9.34E-03 |
| 361 | Retinal detachments and defects | 34 | 7461 | 0.454% | 317 | 33647 | 0.933% | 5.51E-05 |
| 365.1 | Open-angle glaucoma | 173 | 9195 | 1.847% | 57 | 33647 | 0.169% | 8.88E-86 |
| 365.11 | Primary open angle glaucoma | 95 | 9195 | 1.023% | 57 | 33647 | 0.169% | 4.35E-34 |
| 368.2 | Diplopia and disorders of binocular vision | 62 | 9444 | 0.652% | 61 | 34332 | 0.177% | 2.12E-14 |
| 377 | Disorders of optic nerve and visual pathways | 32 | 7639 | 0.417% | 28 | 34026 | 0.082% | 8.68E-12 |
| 378 | Strabismus and other disorders of binocular eye movements | 58 | 9399 | 0.613% | 134 | 34026 | 0.392% | 5.34E-03 |
| 382 | Otalgia | 90 | 9669 | 0.922% | 23 | 34296 | 0.067% | 1.84E-48 |
| 389.4 | Tinnitus | 118 | 9397 | 1.240% | 47 | 34285 | 0.137% | 6.66E-54 |
| 425 | Cardiomyopathy | 1899 | 7211 | 20.845% | 95 | 34369 | 0.276% | 0.00E+00 |
| 425.1 | Primary/intrinsic cardiomyopathies | 1643 | 7211 | 18.557% | 93 | 34369 | 0.270% | 0.00E+00 |
| 425.11 | Hypertrophic obstructive cardiomyopathy | 120 | 7211 | 1.637% | 7 | 34369 | 0.020% | 5.71E-114 |
| 425.12 | Other hypertrophic cardiomyopathy | 65 | 5788 | 1.111% | 5 | 34369 | 0.015% | 7.93E-76 |
| 425.2 | Secondary/extrinsic cardiomyopathies | 629 | 7211 | 8.023% | 1* | 34369* | 0.003%* | 0.00E+00 |
| 426.8 | Other cardiac conduction disorders | 39 | 2431 | 1.579% | 4 | 32071 | 0.012% | 9.99E-98 |
| 426.9 | Cardiac pacemaker/device in situ | 1885 | 3255 | 36.673% | 91 | 32071 | 0.283% | 0 |
| 426.92 | Cardiac defibrillator in situ | 1291 | 3255 | 28.399% | 168* | 33138* | 0.504%* | 0 |
| 427.1 | Paroxysmal tachycardia, unspecified | 1721 | 3255 | 34.586% | 310 | 32071 | 0.957% | 0 |
| 427.12 | Paroxysmal ventricular tachycardia | 1345 | 3255 | 29.239% | 67 | 32071 | 0.208% | 0 |
| 427.6 | Premature beats | 797 | 3255 | 19.669% | 73 | 32071 | 0.227% | 0 |
| 430 | Intracranial hemorrhage | 113 | 8076 | 1.380% | 144 | 33742 | 0.425% | 5.38E-23 |
| 433.6 | Acute, but ill-defined cerebrovascular disease | 136 | 8076 | 1.656% | 308* | 33742* | 0.905%* | 2.93E-09 |
| 442.3 | Aneurysm of artery of lower extremity | 33 | 5772 | 0.568% | 14 | 33943 | 0.041% | 3.06E-26 |
| 444 | Arterial embolism and thrombosis | 107 | 7344 | 1.436% | 71 | 33943 | 0.209% | 3.85E-48 |
| 446 | Polyarteritis nodosa and allied conditions | 46 | 5772 | 0.791% | 77 | 33943 | 0.226% | 1.90E-12 |
| 447 | Other disorders of arteries and arterioles | 590 | 7344 | 7.436% | 101 | 33943 | 0.297% | 0.00E+00 |
| 447.7 | Aortic ectasia | 143 | 7344 | 1.910% | 101* | 33943* | 0.297%* | 7.83E-61 |
| 450 | Noninfectious disorders of lymphatic channels | 86 | 10133 | 0.842% | 70 | 34559 | 0.202% | 1.27E-21 |

| | | | | | | | | |
|---|---|---|---|---|---|---|---|---|
| 451.2 | Phlebitis and thrombophlebitis of lower extremities | 75 | 6650 | 1.115% | 411 | 30719 | 1.320% | 1.95E-01 |
| 454.11 | Varicose veins of lower extremity, symptomtic | 61 | 6650 | 0.909% | 57 | 30719 | 0.185% | 2.80E-21 |
| 458.1 | Orthostatic hypotension | 95 | 7677 | 1.222% | 119 | 32771 | 0.362% | 9.40E-21 |
| 480.12 | Pseudomonal pneumonia | 37 | 6838 | 0.538% | 7 | 33646 | 0.021% | 1.82E-31 |
| 496.2 | Chronic bronchitis | 232 | 7481 | 3.008% | 355 | 30277 | 1.159% | 5.61E-32 |
| 496.3 | Bronchiectasis | 73 | 6094 | 1.184% | 251 | 30277 | 0.822% | 7.07E-03 |
| 499 | Cystic fibrosis | 27 | 8117 | 0.332% | 6 | 34623 | 0.017% | 3.04E-19 |
| 500.2 | Pneumoconiosis | 21 | 6180 | 0.339% | 17 | 33667 | 0.050% | 6.33E-11 |
| 508 | Pulmonary collapse; interstitial and compensatory emphysema | 89 | 7721 | 1.140% | 172 | 33667 | 0.508% | 3.12E-10 |
| 514.1 | Abnormal results of function study of pulmonary system | 23 | 7366 | 0.311% | 3 | 34417 | 0.009% | 3.30E-20 |
| 521.1 | Dental caries | 68 | 9630 | 0.701% | 421 | 33367 | 1.246% | 9.40E-06 |
| 526.4 | Temporomandibular joint disorders | 24 | 7787 | 0.307% | 25 | 33367 | 0.075% | 2.19E-07 |
| 528 | Diseases of the oral soft tissues, excluding lesions specific for gingiva and tongue | 105 | 9774 | 1.063% | 485 | 33956 | 1.408% | 9.59E-03 |
| 536.3 | Gastroparesis | 112 | 9111 | 1.214% | 361* | 31376* | 1.137%* | 5.80E-01 |
| 550.5 | Ventral hernia | 104 | 8901 | 1.155% | 351 | 29721 | 1.167% | 9.69E-01 |
| 555.2 | Ulcerative colitis | 55 | 6400 | 0.852% | 292 | 27446 | 1.053% | 1.68E-01 |
| 558 | Noninfectious gastroenteritis | 146 | 7832 | 1.830% | 1634 | 27446 | 5.619% | 1.79E-44 |
| 560.4 | Other intestinal obstruction | 88 | 7832 | 1.111% | 286 | 27446 | 1.031% | 5.81E-01 |
| 565 | Anal and rectal conditions | 173 | 9351 | 1.816% | 1494 | 32501 | 4.395% | 6.83E-31 |
| 571.6 | Primary biliary cirrhosis | 20 | 6944 | 0.287% | 18 | 33865 | 0.053% | 1.91E-08 |
| 571.81 | Portal hypertension | 50 | 6944 | 0.715% | 34 | 33865 | 0.100% | 2.17E-24 |
| 572 | Ascites (non malignant) | 178 | 8607 | 2.026% | 100 | 33865 | 0.294% | 7.47E-72 |
| 573 | Other disorders of liver | 149 | 8607 | 1.702% | 134 | 33865 | 0.394% | 7.76E-41 |
| 574.2 | Calculus of bile duct | 34 | 7722 | 0.438% | 243 | 32964 | 0.732% | 5.75E-03 |
| 574.3 | Cholecystitis without cholelithiasis | 33 | 7722 | 0.426% | 281 | 32964 | 0.845% | 1.80E-04 |
| 593.2 | Microscopic hematuria | 84 | 7987 | 1.041% | 1392* | 32001* | 4.169%* | 5.53E-42 |
| 596 | Other disorders of bladder | 227 | 9577 | 2.315% | 872 | 33338 | 2.549% | 2.04E-01 |
| 604 | Disorders of penis | 25 | 6680 | 0.373% | 335 | 32616 | 1.017% | 5.85E-07 |
| 618.1 | Prolapse of vaginal walls | 34 | 8036 | 0.421% | 731 | 33489 | 2.136% | 4.13E-25 |
| 681.3 | Cellulitis and abscess of arm/hand | 65 | 8987 | 0.718% | 593 | 33451 | 1.742% | 2.35E-12 |
| 681.6 | Cellulitis and abscess of foot, toe | 22 | 7314 | 0.300% | 593 | 33451 | 1.742% | 3.42E-20 |
| 687.4 | Disturbance of skin sensation | 455 | 8527 | 5.066% | 257 | 34103 | 0.748% | 4.18E-180 |
| 689 | Disorder of skin and subcutaneous tissue NOS | 107 | 9851 | 1.075% | 676 | 33953 | 1.952% | 5.46E-09 |

| | | | | | | | | |
|---|---|---|---|---|---|---|---|---|
| 695.7 | Prurigo and Lichen | 49 | 8821 | 0.552% | 93 | 34001 | 0.273% | 6.78E-05 |
| 698 | Pruritus and related conditions | 193 | 9759 | 1.939% | 74 | 34555 | 0.214% | 1.88E-85 |
| 705.8 | Hyperhidrosis | 120 | 9289 | 1.275% | 63 | 33575 | 0.187% | 4.46E-46 |
| 711 | Arthropathy associated with infections | 22 | 7238 | 0.303% | 41 | 30606 | 0.134% | 2.50E-03 |
| 715 | Other inflammatory spondylopathies | 38 | 7225 | 0.523% | 131 | 30539 | 0.427% | 3.14E-01 |
| 726.2 | Synoviopathy | 21 | 7091 | 0.295% | 38 | 31189 | 0.122% | 1.36E-03 |
| 729 | Other disorders of soft tissues | 59 | 8601 | 0.681% | 661 | 31189 | 2.075% | 4.71E-18 |
| 733.4 | Aseptic necrosis of bone | 54 | 9307 | 0.577% | 43 | 32920 | 0.130% | 4.20E-15 |
| 735.2 | Acquired toe deformities | 74 | 9601 | 0.765% | 494 | 33278 | 1.463% | 1.31E-07 |
| 735.21 | Hammer toe (acquired) | 48 | 9601 | 0.497% | 173 | 33278 | 0.517% | 8.74E-01 |
| 735.3 | Hallux valgus (Bunion) | 58 | 9601 | 0.600% | 679 | 33278 | 2.000% | 7.34E-21 |
| 741 | Symptoms and disorders of the joints | 259 | 9573 | 2.634% | 361 | 34011 | 1.050% | 9.09E-32 |
| 741.2 | Stiffness of joint | 25 | 7761 | 0.321% | 87 | 34011 | 0.255% | 3.71E-01 |
| 742 | Derangement of joint, non-traumatic | 29 | 7761 | 0.372% | 281 | 34011 | 0.819% | 4.26E-05 |
| 742.9 | Other derangement of joint | 25 | 7761 | 0.321% | 240 | 34011 | 0.701% | 1.83E-04 |
| 771.2 | Cramp of limb | 117 | 8991 | 1.285% | 77* | 34406* | 0.223%* | 3.29E-41 |
| 772.1 | Muscular wasting and disuse atrophy | 33 | 7723 | 0.425% | 10 | 34564 | 0.029% | 2.71E-22 |
| 772.2 | Spasm of muscle | 30 | 7723 | 0.387% | 1* | 34564* | 0.003%* | 1.88E-28 |
| 772.3 | Muscle weakness | 41 | 7723 | 0.528% | 55* | 34564* | 0.159%* | 1.42E-09 |
| 800 | Fracture of lower limb | 115 | 9520 | 1.194% | 588 | 32766 | 1.763% | 1.25E-04 |
| 830 | Dislocation | 161 | 9870 | 1.605% | 230 | 32697 | 0.699% | 9.59E-17 |
| 870 | Open wounds of head; neck; and trunk | 140 | 9543 | 1.446% | 336 | 33818 | 0.984% | 1.35E-04 |
| 870.3 | Other open wound of head and face | 27 | 7701 | 0.349% | 263 | 33818 | 0.772% | 7.41E-05 |
| 938.2 | Chronic dermatitis due to solar radiation | 86 | 6976 | 1.218% | 24 | 34217 | 0.070% | 3.65E-64 |
| 962.1 | Adrenal cortical steroids causing adverse effects in therapeutic use | 220 | 7679 | 2.785% | 40* | 31852* | 0.125%* | 3.59E-151 |
| 964 | Poisoning by agents primarily affecting blood constituents | 43 | 6245 | 0.684% | 15 | 31852 | 0.047% | 1.86E-31 |
| 964.1 | Anticoagulants causing adverse effects | 39 | 6245 | 0.621% | 13 | 31852 | 0.041% | 4.13E-29 |
| 965.1 | Opiates and related narcotics causing adverse effects in therapeutic use | 65 | 7679 | 0.839% | 100 | 31852 | 0.313% | 2.00E-10 |
| 990 | Effects radiation NOS | 134 | 9241 | 1.429% | 257 | 34173 | 0.746% | 6.80E-10 |

**Table S6. Comparisons of case prevalence for Phecodes of interest in Penn Medicine Biobank versus UK Biobank.**

List of 139 Phecodes that had associations with 106 different genes at p<E-06 from exome-by-phenome-wide association studies using pLOF-based gene burdens. Phecodes are sorted by increasing Phecode number, and are each labeled with number of cases, number of controls, and case prevalence (*i.e.* cases/(cases+controls)) in Penn Medicine Biobank (PMBB) versus UK Biobank (UKBB). Furthermore, each Phecode is labeled with p-value from chi-square test of independence comparing case prevalence in PMBB versus UKBB. *For Phecodes that were not mapped to from any ICD-10 codes in Phecode Map 1.2b1, corresponding ICD-9 codes according to Phecode Map 1.2 were translated to ICD-10 codes, then mapped back to Phecodes via Phecode Map 1.2b1. If the resulting Phecode was different, then the case and control counts for the new Phecode were listed. **ICD-10 R791, which corresponds to the ICD-9 that maps to Phecode 286.9, does not map to any Phecodes in Phecode Map 1.2b1.

### Table S7

| | PMBB pLOF-Based Gene Burden | | | | UKBB pLOF-Based Gene Burden | | | | |
|---|---|---|---|---|---|---|---|---|---|
| **Gene** | **Phecode** | **Description** | **ß** | **P** | **Phecode** | **Description** | **ß** | **P** | **N** |
| *AMHR2* | 296.1 | Bipolar | 3.84 | 6.84E-07 | 296 | Mood disorders | 2.12 | 3.93E-02 | 29 het |
| | | | | | 296.1 | Bipolar | 2.43 | 1.85E-02 | |
| *BRCA2* | 174 | Breast cancer | 2.23 | 1.72E-07 | 174 | Breast cancer | 1.29 | 2.78E-06 | 170 single het, 1 double het |
| | 174.1 | Breast cancer [female] | 2.23 | 1.88E-07 | 174.1 | Breast cancer [female] | 1.29 | 2.63E-06 | |
| | 174.11 | Malignant neoplasm of female breast | 2.26 | 1.36E-07 | 174.11 | Malignant neoplasm of female breast | 1.30 | 3.88E-06 | |
| *CPT1C* | 447 | Other disorders of arteries and arterioles | 2.22 | 2.01E-07 | 447 | Other disorders of arteries and arterioles | 2.58 | 1.26E-02 | 26 het |
| *CYP2D6* | 965.1 | Opiates and related narcotics causing adverse effects in therapeutic use | 2.74 | 1.50E-09 | 965 | Poisoning by analgesics, antipyretics, and antirheumatics | 2.22 | 4.41E-03 | 14 het |
| *MICALL2* | 681.6 | Cellulitis and abscess of foot, toe | 4.35 | 7.75E-07 | 681.7 | Cellulitis and abscess of trunk | 2.71 | 8.42E-03 | 37 het |
| *MYO1A* | 689 | Disorder of skin and subcutaneous tissue NOS | 2.77 | 2.68E-08 | 689 | Disorder of skin and subcutaneous tissue NOS | 0.90 | 1.63E-02 | 128 single het, 3 double het |
| *OR5H6* | 771.2 | Cramp of limb | 1.71 | 9.59E-07 | 771.1 | Swelling of limb | 1.32 | 9.28E-04 | 44 single het, 7 double het |
| *RGS12* | 250.12 | Type 1 diabetes with renal manifestations | 3.87 | 6.48E-08 | 250.4 | Abnormal glucose | 1.47 | 4.05E-03 | 540 single het, 2 double het |
| | 250.13 | Type 1 diabetes with ophthalmic manifestations | 4.04 | 3.36E-08 | 250.41 | Impaired fasting glucose | 1.85 | 1.16E-02 | |
| *TTN* | 425 | Cardiomyopathy | 0.88 | 7.83E-13 | 425 | Cardiomyopathy | 1.80 | 9.46E-10 | 558 single hets, 18 double het |
| | 425.1 | Primary/intrinsic cardiomyopathies | 0.87 | 1.11E-11 | 425.1 | Primary/intrinsic cardiomyopathies | 1.82 | 5.96E-10 | |
| | 426.9 | Cardiac pacemaker/device in situ | 0.85 | 3.69E-07 | 426 | Cardiac conduction disorders | 0.56 | 3.09E-02 | |
| | 426.92 | Cardiac defibrillator in situ | 0.99 | 1.77E-08 | 426.9 | Cardiac pacemaker/device in situ | 1.19 | 6.97E-03 | |
| | 427.1 | Paroxysmal tachycardia, unspecified | 0.94 | 6.45E-09 | 426.91 | Cardiac pacemaker in situ | 1.32 | 2.85E-03 | |
| | 427.12 | Paroxysmal ventricular tachycardia | 0.96 | 2.18E-08 | 427 | Cardiac dysrhythmias | 0.66 | 1.01E-07 | |
| | 427.6 | Premature beats | 1.11 | 8.90E-09 | 427.2 | Atrial fibrillation and flutter | 0.81 | 7.25E-08 | |
| | | | | | 427.9 | Palpitations | 0.97 | 4.95E-05 | |
| *WDR27* | 695.7 | Prurigo and Lichen | 4.17 | 6.31E-07 | 695.9 | Unspecified erythematous condition | 2.76 | 7.26E-03 | 63 het |

**Table S7: Replication via predicted loss-of-function (pLOF)-based gene burdens in UK Biobank.**

List of significantly replicated associations via gene burdens collapsing rare (MAF ≤ 0.1%) pLOF variants in UK Biobank (UKBB) among 106 genes for which predicted loss-of-function (pLOF)-based gene burdens had p<E-06 from exome-by-phenome-wide association studies in Penn Medicine Biobank (PMBB). Summary statistics are based on an exact logistic regression model adjusted for age, age$^2$, gender, and the first ten principal components of ancestry. Reported are the betas and p-values resulting from running the logistic regression model separately in individuals of European ancestry only, and the number of carriers for each gene burden.

### Table S8

| | PMBB pLOF-Based Gene Burden | | | | UKBB REVEL-Informed Missense-Based Gene Burden | | | | |
|---|---|---|---|---|---|---|---|---|---|
| Gene | Phecode | Description | ß | P | Phecode | Description | ß | P | N |
| *ASPH* | 681.3 | Cellulitis and abscess of arm/hand | 4.30 | 5.39E-10 | 681.7 | Cellulitis and abscess of trunk | 2.43 | 1.74E-02 | 53 het |
| *BRCA1* | 433.6 | Acute, but ill-defined cerebrovascular disease | 3.72 | 3.21E-08 | 433.21 | Cerebral artery occlusion, with cerebral infarction | 1.20 | 8.41E-03 | 422 single het, 2 double het |
| *CFTR* | 496.3 | Bronchiectasis | 3.04 | 4.21E-11 | 496 | Chronic airway obstruction | 0.33 | 2.11E-02 | 1145 single het, 13 double het |
| *EFCAB5* | 618.1 | Prolapse of vaginal walls | 4.29 | 3.19E-08 | 618.5 | Prolapse of vaginal vault after hysterectomy | 4.11 | 2.44E-04 | 18 het |
| *GHDC* | 742 | Derangement of joint, non-traumatic | 4.35 | 2.43E-07 | 742 | Derangement of joint, non-traumatic | 2.33 | 2.66E-02 | 12 het |
| | 742.9 | Other derangement of joint | 4.54 | 1.35E-07 | 742.8 | Articular cartilage disorder | 3.45 | 1.08E-03 | |
| | | | | | 742.9 | Other derangement of joint | 2.49 | 1.79E-02 | |
| *MICALL2* | 681.6 | Cellulitis and abscess of foot, toe | 4.35 | 7.75E-07 | 681.2 | Cellulitis and abscess of face/neck | 2.44 | 1.69E-02 | 46 het |
| *POLN* | 572 | Ascites (non malignant) | 2.59 | 1.55E-07 | 572 | Ascites (non malignant) | 1.51 | 3.00E-03 | 303 single het, 1 double het |
| *PPP1R13L* | 365.11 | Primary open angle glaucoma | 3.12 | 7.29E-07 | 365 | Glaucoma | 2.94 | 1.00E-02 | 6 het |
| *RGS12* | 250.12 | Type 1 diabetes with renal manifestations | 3.87 | 6.48E-08 | 250.11 | Type 1 diabetes with ketoacidosis | 2.65 | 1.04E-02 | 1 hom, 125 het |
| | 250.13 | Type 1 diabetes with ophthalmic manifestations | 4.04 | 3.36E-08 | | | | | |
| *RTKN2* | 458.1 | Orthostatic hypotension | 4.44 | 7.24E-07 | 458 | Hypotension | 1.48 | 4.32E-02 | 36 het |
| | | | | | 458.1 | Orthostatic hypotension | 2.35 | 2.34E-02 | |
| *SH3D21* | 735.3 | Hallux valgus (Bunion) | 3.72 | 2.32E-07 | 735 | Acquired foot deformities | 2.34 | 9.17E-03 | 6 het |
| | | | | | 735.3 | Hallux valgus (Bunion) | 2.69 | 3.01E-03 | |
| *SYT15* | 430 | Intracranial hemorrhage | 2.63 | 1.81E-07 | 430 | Intracranial hemorrhage | 2.27 | 2.82E-02 | 23 het |
| | | | | | 430.1 | Subarachnoid hemorrhage | 3.00 | 3.99E-03 | |
| *TTN* | 427.1 | Paroxysmal tachycardia, unspecified | 0.94 | 6.45E-09 | 427.41 | Ventricular fibrillation and flutter | 1.11 | 8.89E-03 | 1 hom, 2704 single het, 84 double het, 4 triple het |
| | 427.12 | Paroxysmal ventricular tachycardia | 0.96 | 2.18E-08 | | | | | |
| | 427.6 | Premature beats | 1.11 | 8.90E-09 | | | | | |
| *ZNF175* | 389.4 | Tinnitus | 3.35 | 3.24E-10 | 389 | Hearing loss | 2.43 | 2.22E-02 | 10 het |
| | | | | | 389.1 | Sensorineural hearing loss | 5.23 | 3.54E-06 | |

**Table S8: Replication via REVEL-informed missense-based gene burdens in UK Biobank.** List of significantly replicated associations via gene burdens collapsing rare (MAF ≤ 0.1%) predicted deleterious missense (REVEL ≥ 0.5) variants in UK Biobank (UKBB) among 106 genes for which predicted loss-of-function (pLOF)-based gene burdens had p<E-06 from exome-by-phenome-wide association studies in Penn Medicine Biobank (PMBB). Summary statistics are based on an exact logistic regression model adjusted for age, $age^2$, gender, and the first ten principal components of ancestry. Reported are the betas and p-values resulting from running the logistic regression model separately in individuals of European ancestry only, and the number of carriers for each gene burden.

**Table S9**

| PMBB pLOF-Based Gene Burden | | | | | UKBB Single Variants | | | | | | | | | |
|---|---|---|---|---|---|---|---|---|---|---|---|---|---|---|
| ene | Phecode | Description | ß | P | Chr | Position | Ref | Alt | Phecode | Description | ß | P | N | rs |
| *SPH* | 681.3 | Cellulitis and abscess of arm/hand | 4.30 | 5.39E-10 | 8 | 61638365 | TA | T | 681.2 | Cellulitis and abscess of face/neck | 0.73 | 2.24E-02 | 3533 het | rs1461257 |
| *ES5A* | 286.9 | Abnormal coagulation profile | 3.31 | 8.10E-08 | 16 | 55866060 | A | T | 286 | Coagulation defects | 2.37 | 1.18E-03 | 55 het | rs1472908 |
| | | | | | | | | | 286.7 | Other and unspecified coagulation defects | 3.62 | 1.45E-06 | | |
| *FTR* | 480.12 | Pseudomonal pneumonia | 2.99 | 2.27E-07 | 7 | 117531068 | T | C | 480.5 | Bronchopneumonia and lung abscess | 2.93 | 4.50E-03 | 69 het | rs355162 |
| | 496.3 | Bronchiectasis | 3.04 | 4.21E-11 | | | | | 496 | Chronic airway obstruction | 0.99 | 3.66E-02 | | |
| | | | | | | | | | 496.2 | Chronic bronchitis | 1.63 | 6.52E-03 | | |
| | | | | | | | | | 496.21 | Obstructive chronic bronchitis | 1.70 | 4.47E-03 | | |
| | | | | | | | | | 496.3 | Bronchiectasis | 2.01 | 8.33E-04 | | |
| | | | | | 7 | 117559594 | T | G | 496.2 | Chronic bronchitis | 1.30 | 1.30E-02 | 104 het | rs745715 |
| | | | | | | | | | 496.21 | Obstructive chronic bronchitis | 1.37 | 8.91E-03 | | |
| *HD1* | 733.4 | Aseptic necrosis of bone | 2.88 | 2.67E-07 | 11 | 6534014 | G | T | 733 | Other disorders of bone and cartilage | 1.31 | 4.41E-03 | 237 het | rs1809182 |
| *CAB5* | 618.1 | Prolapse of vaginal walls | 4.29 | 3.19E-08 | 17 | 30080205 | T | C | 618 | Genital prolapse | 0.58 | 2.44E-02 | 290 het | rs200371 |
| | | | | | | | | | 618.1 | Prolapse of vaginal walls | 0.85 | 2.77E-03 | | |
| *TF2I* | 574.3 | Cholecystitis without cholelithiasis | 3.88 | 1.24E-08 | 7 | 74752088 | A | G | 574.1 | Cholelithiasis | 0.76 | 4.04E-02 | 109 het | rs782360 |
| | | | | | | | | | 574.2 | Calculus of bile duct | 1.43 | 1.55E-02 | | |
| *QCE* | 528 | Diseases of the oral soft tissues, excluding lesions specific for gingiva and tongue | 2.79 | 3.80E-07 | 7 | 2586272 | TTGT CCCG GAG | T | 528.11 | Stomatitis and mucositis (ulcerative) | 2.14 | 3.70E-02 | 130 het | rs773701 |
| *CBP2* | 772.2 | Spasm of muscle | 3.41 | 2.08E-07 | 13 | 77176630 | TA | T | 772 | Symptoms of the muscles | 0.54 | 4.59E-02 | 11918 het | rs711027 |
| *WIL3* | 574.2 | Calculus of bile duct | 3.81 | 2.13E-08 | 22 | 24759985 | TG | T | 574.3 | Cholecystitis without cholelithiasis | 1.24 | 3.64E-02 | 1 hom, 98 het | rs747731 |
| *TOX2* | 365.1 | Open-angle glaucoma | 1.95 | 3.50E-07 | 4 | 184009160 | G | T | 365.1 | Open-angle glaucoma | 2.00 | 6.19E-03 | 166 het | rs7814698 |
| | | | | | | | | | 365.11 | Primary open angle glaucoma | 2.00 | 6.19E-03 | | |
| *RDN* | 735.2 | Acquired toe deformities | 3.31 | 3.90E-07 | 6 | 123464887 | GA | G | 735.21 | Hammer toe (acquired) | 1.40 | 1.82E-02 | 169 het | rs201431 |
| | | | | | 6 | 123464887 | G | GA | 735.1 | Flat foot | 1.31 | 9.53E-03 | 2347 het | rs770690 |
| *TTN* | 425 | Cardiomyopathy | 0.88 | 7.83E-13 | 2 | 178566349 | C | G | 427.41 | Ventricular fibrillation and flutter | 1.26 | 4.11E-02 | 6 hom, 1330 het | rs5630721 |
| | 425.1 | Primary/intrinsic cardiomyopathies | 0.87 | 1.11E-11 | 2 | 178573986 | A | G | 426.23 | Second degree AV block | 2.10 | 4.29E-02 | 131 het | rs563992 |

| | | | | | | | | | | | | | |
|---|---|---|---|---|---|---|---|---|---|---|---|---|---|
| 426.9 | Cardiac pacemaker/device in situ | 0.85 | 3.69E-07 | 2 | 178575436 | C | G | 427.5 | Arrhythmia (cardiac) NOS | 0.95 | 2.19E-02 | 12 hom, 989 het | rs5580113 |
| 426.92 | Cardiac defibrillator in situ | 0.99 | 1.77E-08 | 2 | 178576311 | C | T | 426.24 | Atrioventricular block, complete | 2.31 | 3.35E-02 | 77 het | rs2010439 |
| 427.1 | Paroxysmal tachycardia, unspecified | 0.94 | 6.45E-09 | | | | | 426.4 | Anomalous atrioventricular excitation | 2.82 | 6.41E-03 | | |
| 427.12 | Paroxysmal ventricular tachycardia | 0.96 | 2.18E-08 | | | | | 427.2 | Atrial fibrillation and flutter | 0.87 | 2.55E-02 | | |
| 427.6 | Premature beats | 1.11 | 8.90E-09 | 2 | 178577205 | G | A | 427.6 | Premature beats | 1.21 | 3.76E-02 | 2 hom, 419 het | rs5598049 |
| | | | | 2 | 178584852 | C | T | 427.1 | Paroxysmal tachycardia, unspecified | 1.50 | 3.91E-02 | 51 het | rs1506619 |
| | | | | | | | | 427.11 | Paroxysmal supraventricular tachycardia | 1.74 | 1.70E-02 | | |
| | | | | | | | | 427.6 | Premature beats | 2.27 | 2.66E-02 | | |
| | | | | 2 | 178614561 | A | G | 427.1 | Paroxysmal tachycardia, unspecified | 1.49 | 4.00E-02 | 54 het | rs7267724 |
| | | | | | | | | 427.12 | Paroxysmal ventricular tachycardia | 2.26 | 2.67E-02 | | |
| | | | | | | | | 427.4 | Cardiac arrest and ventricular fibrillation | 2.10 | 4.04E-02 | | |
| | | | | | | | | 427.41 | Ventricular fibrillation and flutter | 3.44 | 1.02E-03 | | |
| | | | | 2 | 178653276 | A | G | 426 | Cardiac conduction disorders | 0.35 | 7.60E-03 | 72 hom, 2947 het | rs256284 |
| | | | | | | | | 426.3 | Bundle branch block | 0.41 | 1.34E-02 | | |
| | | | | | | | | 426.32 | Left bundle branch block | 0.44 | 4.74E-02 | | |
| | | | | | | | | 427.2 | Atrial fibrillation and flutter | -0.23 | 2.67E-02 | | |
| | | | | 2 | 178654748 | C | T | 426.23 | Second degree AV block | 2.29 | 2.76E-02 | 127 het | rs7554153 |
| | | | | 2 | 178662368 | G | A | 426.24 | Atrioventricular block, complete | 0.77 | 4.31E-02 | 285 hom, 1087 het | rs2014745 |
| | | | | 2 | 178675734 | G | T | 427.42 | Cardiac arrest | 1.21 | 4.06E-02 | 1 hom, 440 het | rs1824287 |
| | | | | 2 | 178698916 | TAA | T | 426.91 | Cardiac pacemaker in situ | 0.69 | 4.22E-02 | 2336 het | rs7743292 |
| | | | | 2 | 178698916 | TAAA | T | 425 | Cardiomyopathy | 1.38 | 1.95E-02 | 283 het | rs7458315 |
| | | | | | | | | 425.1 | Primary/intrinsic cardiomyopathies | 1.41 | 1.76E-02 | | |
| | | | | 2 | 178698916 | T | TA | 426 | Cardiac conduction disorders | 0.20 | 4.65E-02 | 4 hom, 8727 het | rs7691785 |
| | | | | | | | | 426.32 | Left bundle branch block | 0.36 | 3.38E-02 | | |
| | | | | | | | | 427.2 | Atrial fibrillation and flutter | -0.18 | 7.53E-03 | | |
| | | | | 2 | 178722403 | C | G | 427 | Cardiac dysrhythmias | 0.15 | 2.82E-04 | 880 hom, 8874 het | rs1269316 |
| | | | | | | | | 427.2 | Atrial fibrillation and flutter | 0.15 | 4.57E-03 | | |

| | | | | | | | | | | | | | | |
|---|---|---|---|---|---|---|---|---|---|---|---|---|---|---|
| | | | | | | | | | 427.5 | Arrhythmia (cardiac) NOS | 0.43 | 1.91E-02 | | |
| | | | | | 2 | 178733826 | T | G | 427.42 | Cardiac arrest | 1.61 | 2.68E-02 | 224 het | rs7264893 |
| | | | | | 2 | 178779048 | C | T | 425 | Cardiomyopathy | 1.41 | 5.64E-03 | 1 hom, 396 het | rs3602185 |
| | | | | | | | | | 425.1 | Primary/intrinsic cardiomyopathies | 1.43 | 4.96E-03 | | |
| *DR27* | 695.7 | Prurigo and Lichen | 4.17 | 6.31E-07 | 6 | 169636370 | C | A | 695 | Erythematous conditions | 0.97 | 3.17E-02 | 2 hom, 262 het | rs1116566 |
| | | | | | | | | | 695.7 | Prurigo and Lichen | 1.43 | 1.42E-02 | | |
| *DR87* | 550.5 | Ventral hernia | 2.12 | 1.70E-07 | 19 | 37888807 | G | A | 550.3 | Femoral hernia | 1.71 | 1.89E-02 | 149 het | rs1897043 |

**Table S9. Replication via univariate association studies in UK Biobank.**

List of significantly replicated associations via univariate analyses of low-frequency to common (MAF > 0.1%) predicted loss-of-function (pLOF) or predicted deleterious missense (REVEL ≥ 0.5) variants UK Biobank (UKBB) among 106 genes for which predicted loss-of-function (pLOF)-based gene burdens had p<E-06 from exome-by-phenome-wide association studies in Penn Medicine Biobank (PMBB). Summary statistics are based on an exact logistic regression model adjusted for age, age$^2$, gender, and the first ten principal components of ancestry. Reported are the betas and p-values resulting from running the logistic regression model separately in individuals of European ancestry only, and the number of carriers for each gene burden or single variant. Additionally, single variants are annotated by their genomic location according to GRCh38/hg38, as well as their rs identification codes.

**Table S10**

| Gene | Echo parameter | Carrier Median (IQR) N | Non-Carrier Median (IQR) N | ß | p |
|---|---|---|---|---|---|
| *TTN* | LVOT velocity time integral, median (cm) | 18.1 (15.7, 21.6)<br>77 | 21.0 (17.4, 24.7)<br>2196 | -1.96 | 5.34E-04 |
| | Left ventricular ejection fraction (LVEF), minimum | 50.0 (35.0, 60.0)<br>117 | 60.0 (45.0, 60.0)<br>3252 | -5.66 | 0.0103 |
| *MYBPC3* | LVOT velocity time integral, median (cm) | 26.3 (22.4, 28.6)<br>11 | 20.8 (17.3, 24.6)<br>2262 | 4.75 | 2.83E-03 |
| | Left ventricular ejection fraction (LVEF), maximum | 65.0 (65.0, 70.0)<br>14 | 65.0 (55.0, 70.0)<br>3202 | 4.66 | 4.02E-04 |
| *BBS10* | LVOT stroke volume (mL), maximum | 121.4 (114.4, 128.4)<br>2 | 76.3 (60.9, 93.6)<br>1314 | 43.97 | 1.91E-03 |
| | Mitral E/A ratio, median | 1.6 (1.2, 1.8)<br>12 | 1.2 (0.8, 1.6)<br>2760 | 0.23 | 0.0487 |

**Table S10. Echocardiographic measurements of cardiomyopathy-associated gene burdens in Penn Medicine Biobank.**

Comparison of representative echocardiography parameters for cardiac size and functionality between heterozygous carriers of candidate gene burdens and individuals in Penn Medicine Biobank not carrying any of the respective gene's pLOF variants with echo data available. Data is represented as beta and p-value for robust linear regression adjusted for age, gender, and the first four principal components of genetic ancestry. Analyses were limited to European ancestry only.
